# Reconstruction of a rare Δ10 desaturation pathway enables production of yellow peach moth pheromone precursors in *Camelina sativa*

**DOI:** 10.64898/2026.10.20.760337

**Authors:** Xiao-Qing Hou, Hong-Lei Wang, Panpan He, Yujie Jin, Xia Gao, Jean-Marc Lassance, Christer Löfstedt, Bao-Jian Ding

**Affiliations:** Xianghu Laboratory, Hangzhou, China; Department of Biology, Lund University, Lund, Sweden; Gembloux Agro-BioTech, University of Liège, Gembloux, Belgium

**Keywords:** pheromone biosynthetic pathway, heterologous expression, functional assays, bioproduction, metabolic engineering, pest control

## Abstract

Sex pheromones are environmentally friendly alternatives to traditional insecticides for pest control. The high synthesis cost, however, has restricted large-scale application. Plant-based biofactories provide a promising, sustainable route for cost-effective production that could facilitate the broader use of pheromone-based pest control. Mating disruption of the yellow peach moth, *Conogethes punctiferalis*–a serious pest of fruit trees and field crops in East Asia–can be achieved using (*E*)-10-hexadecenal and (*Z*)-10-hexadecenal, two components of its sex pheromone. To enable rational reconstruction of the biosynthetic pathway towards these unusual fatty acid derivatives in heterologous production systems, we first elucidated their biosynthesis by combining *in vivo* labeling experiments, transcriptomics and functional characterization of candidate desaturase and reductase genes using *Nicotiana benthamiana* and yeast expression systems. We identified a rare Δ10 fatty acyl desaturase that catalyzes the production of both (*E*)- 10- and (*Z*)-10-hexadecenoic acids, along with a downstream fatty acyl reductase with higher activity on the *E* isomer. We further introduced this Δ10 desaturase together with an acyl-ACP thioesterase-1 into *Camelina sativa* to redirect seed lipid metabolism towards the accumulation of the pheromone precursor fatty acids. After four generations of selection, we obtained stably inherited Camelina lines accumulating up to 4.7% of target fatty acids in total seed lipids, with individual seeds containing up to 8.3 μg, without significant effects on plant growth or seed viability. These findings demonstrate how elucidating moth pheromone biosynthetic pathways can support the development of sustainable biotechnological production systems for pheromone-based pest management.

## 1 Introduction

The yellow peach moth, *Conogethes punctiferalis* (Lepidoptera: Crambidae), is a destructive pest of agricultural significance across Asia, causing losses to multiple crops. This polyphagous insect infests a wide range of host plants, including corn, sunflower, chestnut, peach, and other economically important crops (Thyagaraj et al. 2003). The larvae bore into fruits, seeds, and ears, damaging marketable tissues while remaining concealed within the host plants, which limits the efficacy of conventional chemical pesticide treatments against such internal feeders. This feeding behavior, coupled with increasing concerns over pesticide resistance, environmental pollution and risks to human health, has created an urgent demand for developing more sustainable and species-specific pest management strategies. As an alternative to conventional pesticides, sex pheromone-based approaches, including monitoring, mass trapping and mating disruption have shown promise for the sustainable management of *C. punctiferalis* populations while significantly reducing pesticide usage (Chen et al. 2022; Dong et al. 2024; Kim et al. 2024). For instance, in hickory nut and chestnut orchards, mating disruption achieved >90% male catch inhibition, and significantly reduced its population and the damages on fruits (Dong et al. 2024; Kim et al. 2024). Nevertheless, the high cost and reliance on petrochemicals associated with conventional pheromone synthesis remain major barriers to large-scale deployment of pheromone-based pest management in crop protection, highlighting the need for cost-effective, safe, and efficient platforms for insect pheromone production.

The sex pheromone of *C. punctiferalis* consists of (*E*)-10-hexadecenal (E10-16:Ald) as the major component, along with (*Z*)-10-hexadecenal (Z10-16:Ald) as a minor component (Konno et al. 1982), both classified as Type I pheromones (Ando et al. 2004; Löfstedt et al. 2016). Although additional Type II pheromone compounds, such as (*Z*)-9-heptacosene and (3*Z*,6*Z*,9*Z*)-tricosatriene, have been reported to enhance trap catches (Xiao et al. 2012; Nam et al. 2022), these compounds are not required for effictive mating disruption (Kim et al. 2024). Accordingly, the present study focuses on the major Type I pheromone components. In female moths, Type I pheromone biosynthesis typically begins from saturated fatty acids, such as myristic acid and palmitic acid, and proceeds through sequential reactions including desaturation, chain shortening or elongation, reduction, and functional group modification (Tillman et al. 1999; Ma and Ramaswamy 2003). Among these reactions, desaturation plays a central role in pheromone diversification among moth species. Fatty acyl-CoA desaturases (FADs) introduce double bonds at specific positions and with defined geometries along the fatty acid chain, thereby generation much of the structural diversity of unsaturated pheromone precursors (Bjostad and Roelofs 1983; Knipple et al. 2002; Löfstedt et al. 2016).

Palmitic acid, a key substrate for desaturases in Type I sex pheromone biosynthesis, naturally accumulates in the seeds of oil crops, and its abundance can be further increased by redirecting acyl-ACP metabolism towards C16 fatty acids through manipulation acyl-ACP thioesterase activity. In particular, FATB-type thioesterases can be exploited to increae the availability of C16 fatty acid substrates, providing a sustainable metabolic background for the production of C16 pheromone precursors. By introducing insect-derived fatty acyl desaturases into the seed oil biosynthesis pathway, it is feasible to channel the endogenous fatty acid flux toward the desired pheromone precursors (Löfstedt and Xia 2021). For example, *Camelina sativa*, also known as gold-of-pleasure, or false flax, was successfully engineered to produce (*Z*)-11-hexadecenoic acid (Z11-16:COOH), which serves as the immediate precursor for the aldehyde, alcohol, and acetate pheromone components of a number of moth species. Importantly, the plant-derived pheromones showed similar biological activity to synthetic ones in field trapping and mating disruption tests for *Plutella xylostella* (Wang et al. 2022). Similarly, Camelina-derived (*E8*,*E*10)-dodecadienol also exhibited comparable bioactivity in field trapping and electrophysiological trials (Xia et al. 2021).

The structural diversity of moth pheromones is partly generated by the functional diversitfication of fatty acyl desaturases (FADs), which exhibit remarkable variation in regioselectivity, stereoselectivity, and substrate specificity. To date, desaturases introducing double bonds at several positions, including Δ5, Δ6, Δ9, Δ10, Δ11, Δ14, as well as terminal positions, have been functionally characterized in moths (Foster and Roelofs 1996; Wang et al. 2010; Knipple et al. 1998; Hao et al. 2002; Roelofs et al. 2002; Liu et al. 2002; 2004; Ding et al. 2011; Hagström et al. 2014; Xia 2020; Lassance et al. 2021). Among these activities, Δ10 desaturation appears to be relatively uncommon. Notably, only a single moth Δ10 desaturase has been functionally characterized so far, from the New Zealand leafroller *Planotortrix octo* (Lepidoptera: Tortricidae). This FAD belongs to the Lepidoptera-specific Δ11 desaturase lineage and converts palmitic acid to (*Z*)-10-hexadecenoic acid exclusively (Hao et al. 2002). To our knowledge, a moth desaturase capable of producing the corresponding *trans* isomer (i.e. (*E*)-10-hexadecenoic acid) has not been reported.

Downstream of desaturation, fatty acyl reductases (FARs) convert unsaturated acyl intermediates into fatty alcohols, which may serve directly as sex pheromone components or be further converted into aldehydes or acetate esters (Jurenka 2021). The functionally characterized FARs involved in moth sex pheromone biosynthesis belong to a Lepidoptera-specific pheromone gland FAR (pgFAR) lineage (Lassance et al. 2010). These enzymes can exhibit pronounced substrate selectivity with respect to acyl chain length, double bond position, and double bond geometry, thereby providing an additional layer of specificity in shaping pheromone composition (Liénard et al. 2010; Lassance et al. 2013; Dou et al. 2020; Hagström et al. 2012; Carot-Sans et al. 2015; Ding and Löfstedt 2015).

In the present study, we elucidated the biosynthetic pathway of the sex pheromone components produced by female *C. punctiferalis* through *in vivo* labeling experiments using a deuterium-labeled fatty acid precursor. Combining transcriptomics with functional assays in *Nicotiana benthamiana* and yeast, we identified a Δ10 desaturase and a fatty acyl reductase involved in the pheromone biosynthesis pathway. Finally, we successfully reconstituted a minimal pathway for Δ10 pheromone precursor production in *C. sativa* by introducing the *C. punctiferalis* Δ10 desaturase together with an acyl-ACP thioesterase, resulting in the accumulation of the target fatty acids in transgenic seeds. Our study extends the scope of plant-based pheromone production to unusual Δ10 fatty acid derivatives and expands the enzymatic toolkit for engineering specialty fatty acids in oilseed crops.

## 2 Results

### 2.1 Biosynthetic Pathway Elucidation

Saturated and monounsaturated C16 aldehydes were identified in pheromone gland extracts of *C. punctiferalis* (Figure 1A). The monounsaturated adehydes were poorly resolved chromatographically, appearing as a major peak with a tailing shoulder. Subsequent dimethyl disulfide (DMDS) derivatization identified the peak as a mixture of E10- and Z10-16:Ald, with the *Z* isomer representing approximately 4.5% of the mixture (Figure S1). This proportion is consistent with the 100:8 E10-16:Ald:Z10-16:Ald blend ratio previously reported as optimal in field trials with Chinese populations of *C. punctiferalis* (Liu et al., 1994). Analysis of fatty acids as their methyl esters identified palmitic acid and the corresponding E/Z10-16:COOH (Figure 1B, Figure S2). E10-16:Me was the predominant isomer, while Z10-16:Me was present as a substantial shoulder, suggesting a higher relative abundance of the Z isomer in precursors than observed for the corresponding aldehydes. In addition, several ubiquitous fatty acids, including stearic, oleic, linoleic, and linolenic acids, were also detected in *C. punctiferalis* pheromone glands (Figure 1B).

**Figure 1.**
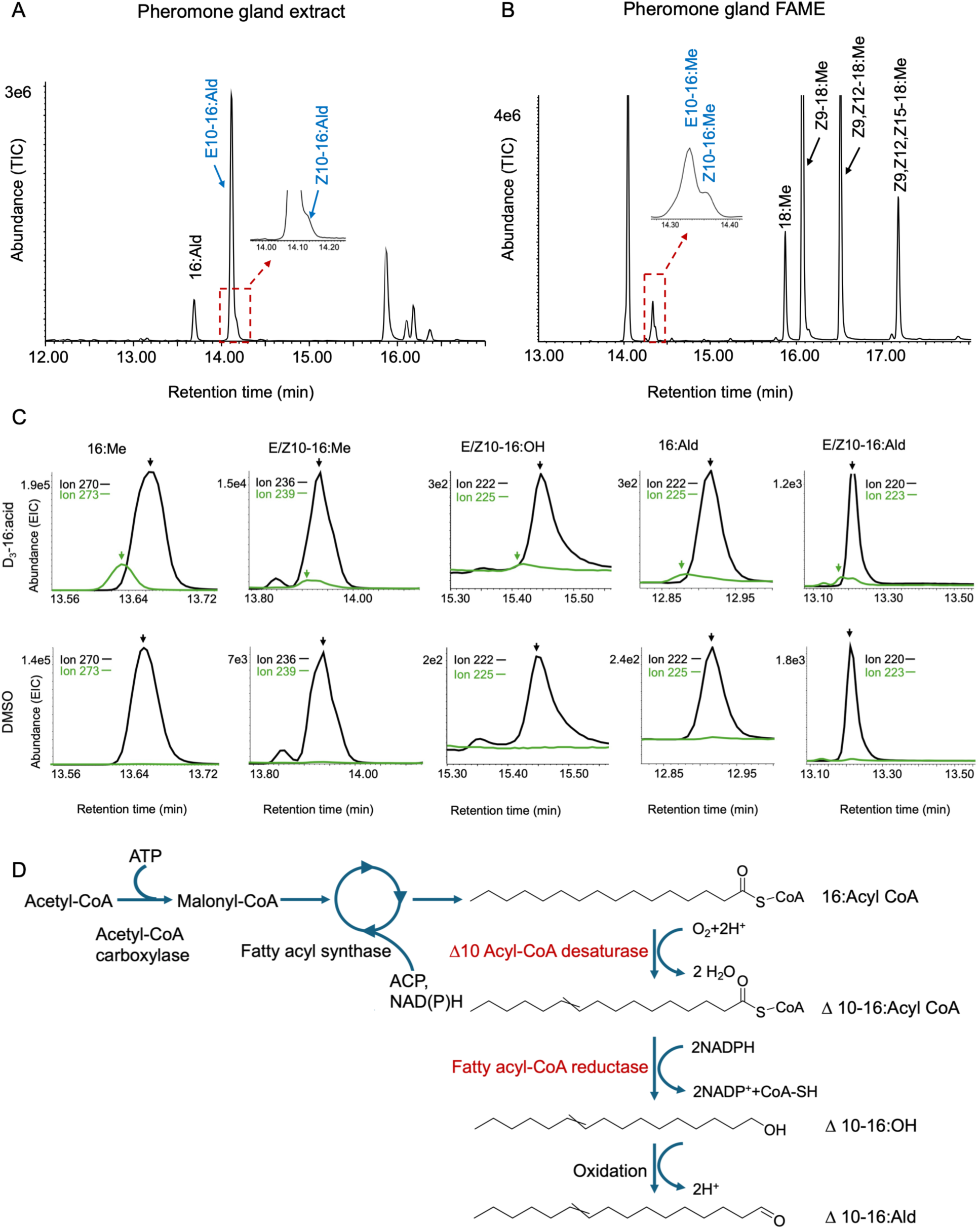
Identification of the biosynthetic pathway of E/Z10-16:Ald in *C. punctiferalis*. (A) Representative chromatogram of pheromone gland heptane extract. The inset shows an enlarged view of the partially resolved E10- and Z10-16:Ald peaks. (B) Representative chromatogram of pheromone gland fatty acids analyzed as fatty acid methyl esters (FAME). The inset shows an enlarged view of the partially resolved E10- and Z10-16:Me peaks. TIC: total ion current. (C) Incorporation of deuterium from [16,16,16– ^2^H_3_] hexadecanoic acid (D_3_–16:COOH) into fatty acid precursors, alcohol intermediates, and aldehydes in *C. punctiferalis* pheromone glands. Representative selected-ion-monitoring traces are shown for glands treated with D_3_–16:COOH (top) and DMSO controls (bottom). Native 16:Me, E/Z10-16:Me, E/Z10-16:OH, 16:Ald, and E/Z10-16:Ald were monitored using ions with *m/z* 270, 236, 222, 222 and 220, respectively, and the corresponding D_3_-labeled compounds were monitored, using ions with *m/z* 273, 239, 225, 225 and 223, respectively. Black and green arrowheads indicate the native and corresponding D_3_-labeled molecules. EIC, extracted ion chromatograms. (D) Proposed biosynthetic pathway of E/Z10-16:Ald in *C. punctiferalis*. The desaturation and reduction steps investigated in this study are highlighted in red.

Deuterium-labeled D_3_–16:COOH was topically applied to the pheromone glands of *C. punctiferalis* to trace the biosynthetic pathway of the major sex pheromone components. Both native and labeled E/Z10:Me, E/Z10-16:OH, 16:Ald, and E/Z10-16:Ald were detected, demonstrating that the incorporation of deuterium atoms from D_3_–16:COOH into the unsaturated pheromone precursors, the corresponding alcohol intermediates, and the aldehyde pheromone components (Figure 1C). In contrast, although native 16:OH was detected, no incorporation of deuterium into the latter was observed. These labeling results support a biosynthetic pathway in which palmitic acid is desaturated at the Δ10 position to produce E10- and Z10-16:COOH, which are subsequently reduced to the corresponding alcohols and oxidized to the aldehyde pheromone components (Figure 1D).

### 2.2 Identification of Desaturase and Reductase Candidates of Sex Pheromone Biosynthetic Pathway from Transcriptomes

We reconstructed the *C. punctiferalis* transcriptome using female pheromone glands, female antennae, female abdomens, and male abdominal tip tissues to create a database allowing comprehensive identification of FADs and FARs potentially involved in sex pheromone biosynthesis. Approximately32 Gb of sequence data were generated, corresponding to 50 million 100-bp paired-end reads per sample. Clean reads from all four samples were co-assembled into 121,257 non-redundant transcripts, with a mean length of 1,418 bp and an N50 of 2,612 bp. In total, 9 FAD and 16 FAR candidates were identified.

We performed phylogenetic analyses of FADs and FARs using predicted amino acid sequences from *C. punctiferalis* and other moth species. The 9 FAD candidates were distributed across the gene tree, with two clustering within the Δ10/Δ11 clade (Figure 2A). Among these, *Cpun_LATN* was highly expressed in the female pheromone glands, (Figure 2A), making it a strong candidate for involvement in the pheromone biosynthesis pathway. In the FAR phylogeny, only Cpun_3376 and Cpun_11842 clustered within the lineage containing functionally characterized pheromone biosynthetic FAR genes (pgFAR; Figure 2B). Cpun_3376 was prioritized for functional characterization based on its high expression value in female pheromone glands (Figure 2B).

**Figure 2.**
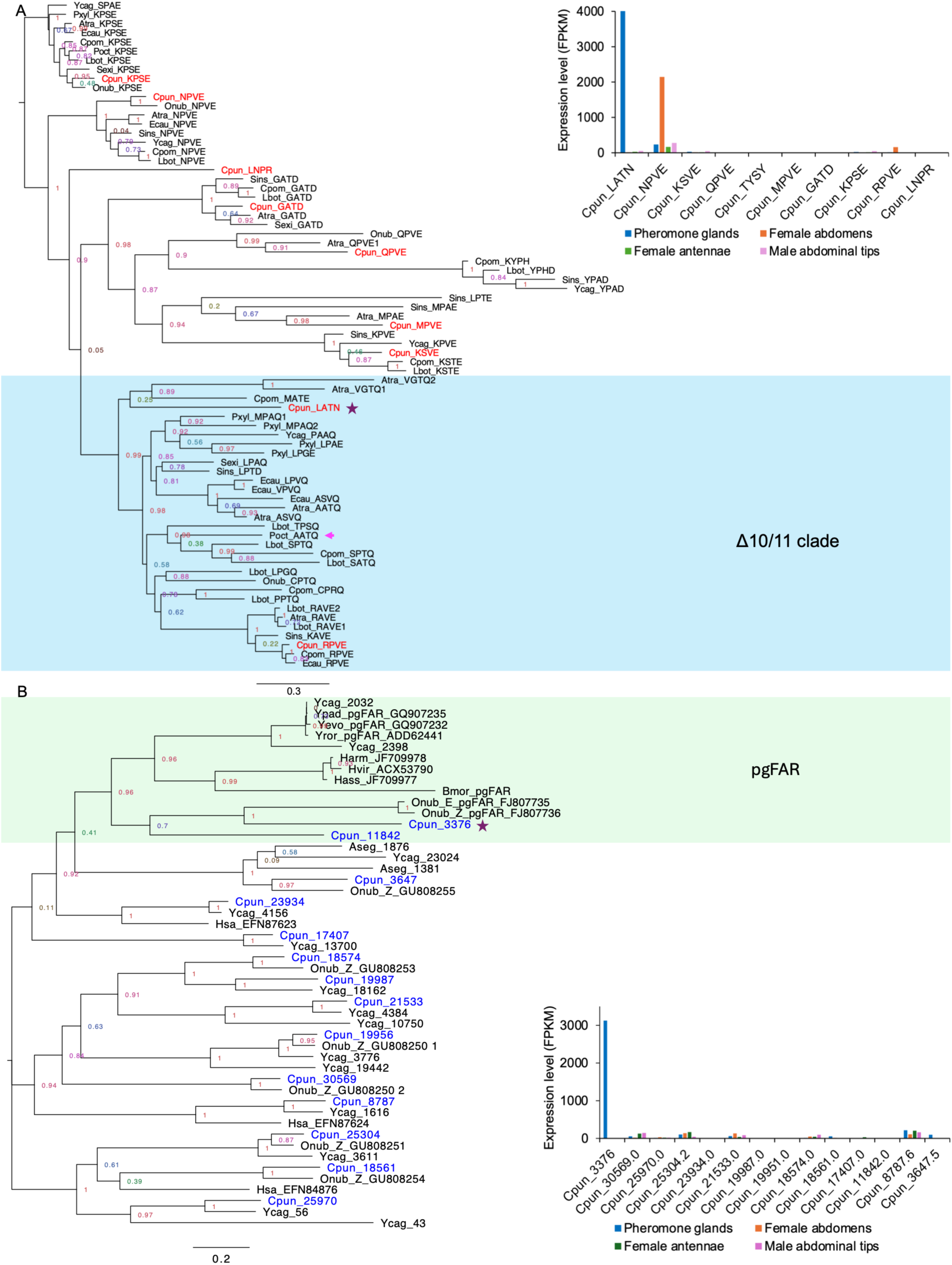
Phylogenetic relationships and tissue expression of candidate fatty acyl desaturases and reductases from *C. punctiferalis*. (A) Maximum-likelihood phylogeny of fatty acyl desaturases (FADs) built based on predicted amino acid sequences of from Lepidoptera. *C. punctiferalis* FADs are shown in red, and the candidate pheromone desaturase is indicated by a plum-colored star. The functionally characterised *Δ*10 desaturase Poct_AATQ is indicated by a pink-colored arrowhead. The Δ10/Δ11 desaturase clade is highlighted in light blue. The inset in the upper-right corner shows expression estimates for *C. punctiferalis* FADs across tissues. (B) Maximum-likelihood phylogeny of fatty acyl reductases (FARs) based on predicted amino acid sequences from Lepidoptera. *C. punctiferalis* FARs are shown in blue, and the candidate pheromone gland reductase is indicated by a plum-colored star. The pgFAR clade is highlighted in light green. The inset in the bottom-right corner shows expression estimates for *C. punctiferalis* FARs across tissues. Expression is reported as fragments per kilobase of transcript per million reads (FPKM). Species abbreviations are defined in Table S1.

### 2.3 Functional characterizaton of Cpun_LATN and Cpun_3376

We next sought to characterize the activities of the candidate FAD and FAR. GC/MS analysis revealed that *N. benthamiana* co-expressing *Cpun_LATN* and *TE16* produced E10-16:COOH and Z10-16:COOH at an approximately 3:1 ratio, whereas neither product was detected in plants expressing *TE16* alone (Figure 3A, 3B; Table S2). Expressing the FAD candidate *Cpun_LATN* in the Δ9 desaturase-deficient (Δ*ole1*) yeast in the presence of oleic acid and supplemented with the substrates 16:Me, 14:Me, and 12:Me did not result in detectable desaturation product (Figure S3). Together, these results demonstrate that *Cpun_LATN* encodes a Δ10 desaturase capable of producing both *E* and *Z* isomers *in planta*.

**Figure 3.**
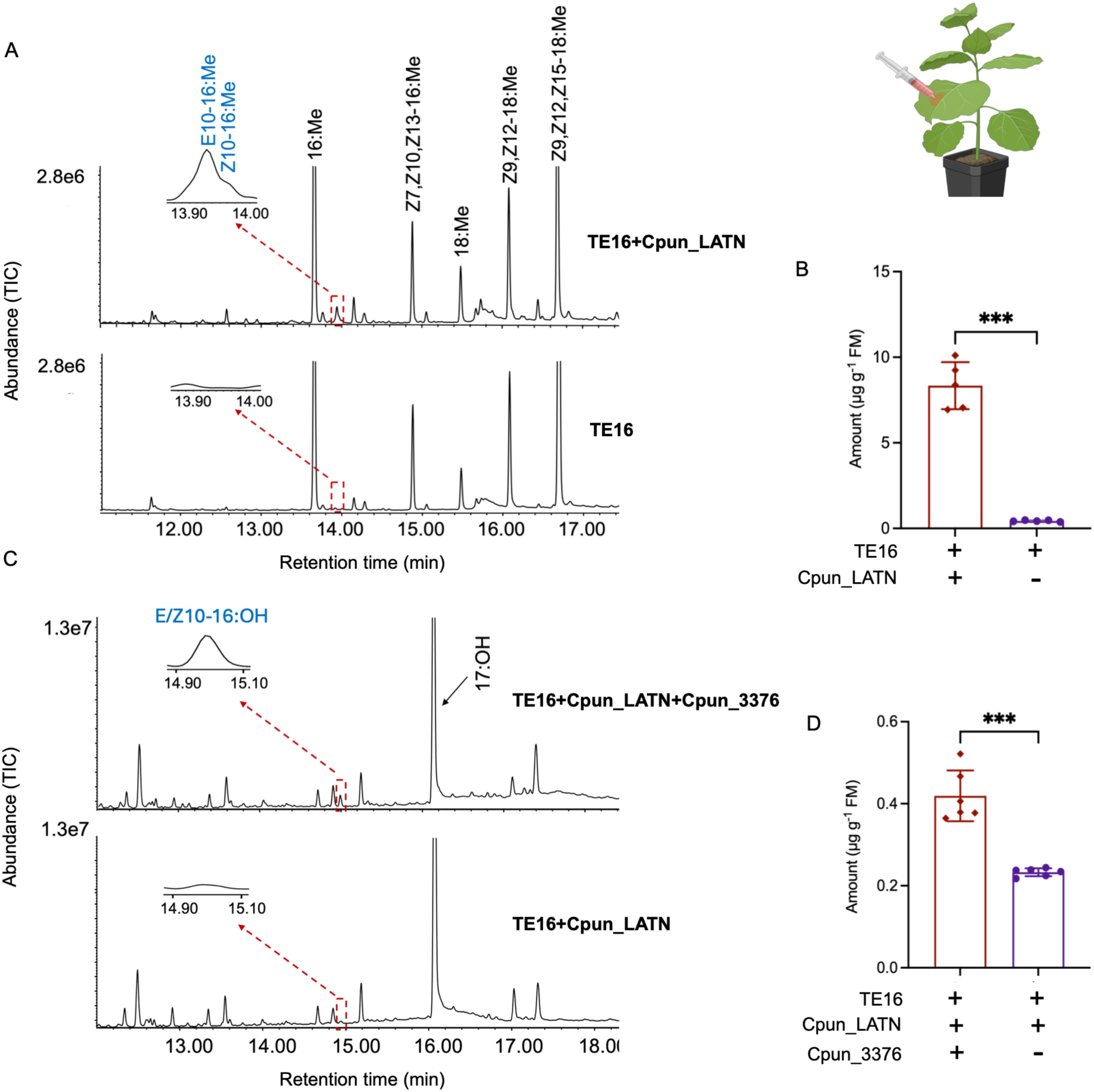
Heterologous expression of fatty acyl desaturase and reductase candidates from *C. punctiferalis* in *N. benthamiana*. (A) Representative chromatogram of fatty acid methyl ester profiles of plant leaves expressing *TE16*/*Cpun_LATN* and *TE16* control. (B) Quantification of E/Z10-16:Me in *N. benthamiana* leaves expressing above the constructs (n=5). (C) Representative chromatogram of fatty alcohol profiles of plant leaves expressing *TE16*/*Cpun_LATN*/*Cpun_3376* and *TE16*/*Cpun_LATN* control. (D) Quantification of E/Z10-16:OH in *N. benthamiana* leaves expressing the above constructs (n=5). The target compounds produced from the introduced genes are labeled in blue. TIC: total ion current. P values are based on t-tests: ***, P < 0.001. Error bars indicate the SEM.

Co-expression of *Cpun_3376*, *Cpun_LATN* and *TE16* in *N. benthamiana* resulted in the production of E/Z10-16:OH, although only at trace levels, preventing reliable quantification of the individual isomers and determination of the E/Z ratio (Figure 3C, 3D; Table S2). The *N. benthamiana* infiltrated with only Cpun_LATN and TE16 showed a tiny peak eluting at close retention time, however, the corresponding mass spectrum does not support it an alcohol compound (Figure S4).

To assess the substrate selectivity of Cpun_3376, yeast expressing the candidate FAR were separately supplied with E10-16:Me or Z10-16:Me substrates, with an unsupplemented culture included as a control. In all three treatments, 16:OH was detected, indicating that Cpun_3376 can reduce endogenous saturated C16 substrate (Figure 4A). Supplementation with E10–16:Me resulted in the production of E10–16:OH, whereas cultures supplied with Z10–16:Me produced only trace amount of Z10–16:OH (Figure 4A, 4B, 4C; Table S3). A small amount of E10–16:OH was also detected in cultures supplied with Z10–16:Me, which can be attributed to the presence of a minor proportion of the E isomer in the Z10-16:Me substrate (Figure S5). No fatty alcohols were detected in yeast transformed with the empty vector, irrespective of substrate supplementation (Figure 4A, 4C; Table S3). Together, these results demonstrate that Cpun_3376 is an active FAR capable of reducing both saturated and Δ10-unsaturated C16 substrates, with a pronounced preference for E10-16 over Z10-16 substrates.

**Figure 4.**
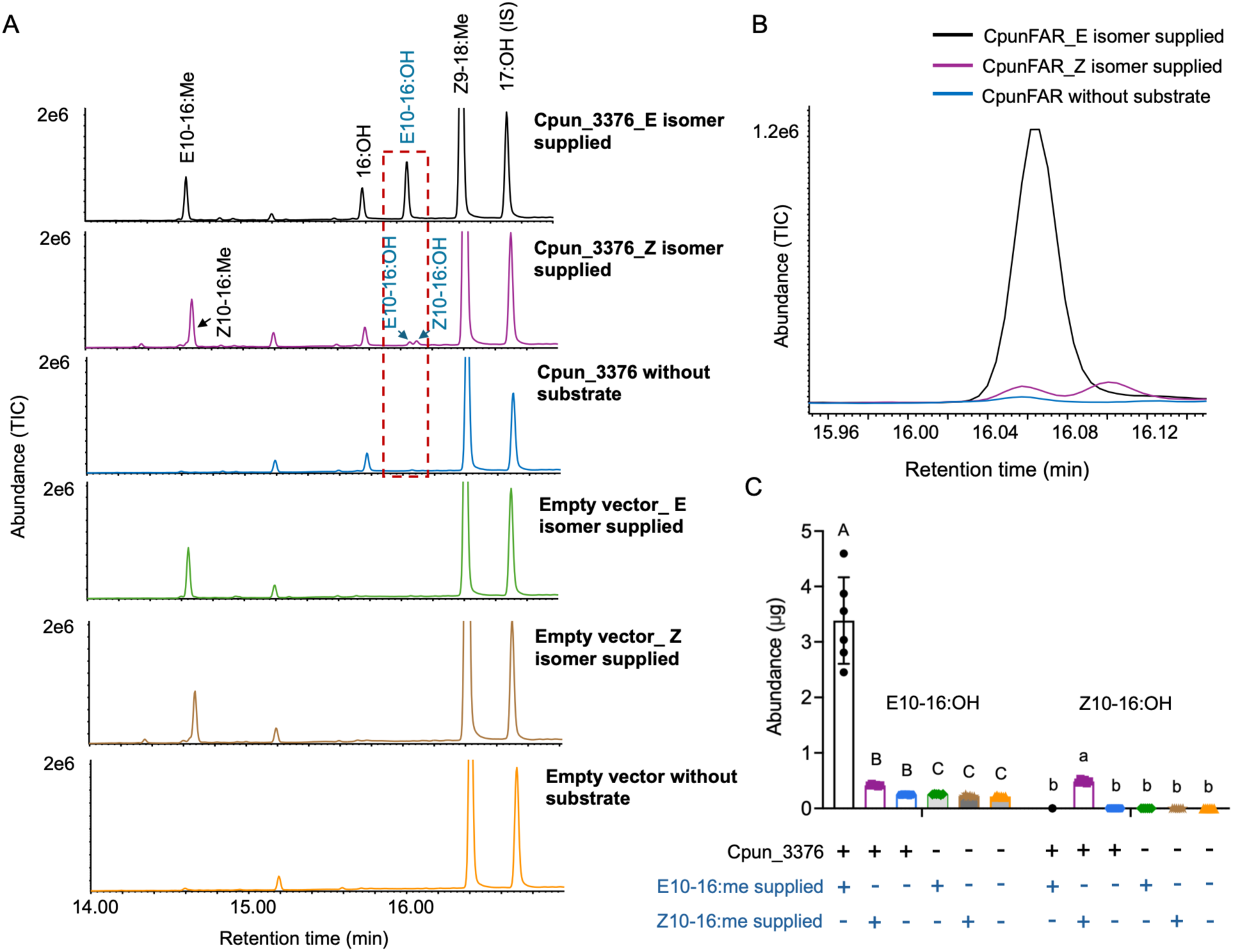
Functional characterization and substrate selectivity of Cpun_3376 in the Δ*ole1* yeast. (A) Representative chromatogram of yeast culture extracts from *ole1* yeast expressing *Cpun_3376* or carrying the empty vector, supplemented with E10-16:Me, Z10-16:Me, or no exogenous substrate. (B) Enlarged and overlaid chromatograms showing Δ10-unsaturated fatty alcohol region for *Cpun_3376*-expressing yeast supplemented with E10-16:Me, Z10-16:Me, or no substrate. TIC: total ion current. (C) Quantification of E10-16:OH and Z10-16:OH in yeast cultures under the different treatments (n=6). Significant differences among treatments are indicated by uppercase and lowercase letters for E10-16:OH and Z10-16:OH, respectively (one-way ANOVA followed by Tukey’s HSD test). Error bars represent SEM.

### 2.4 Bioproduction of E/Z10-16:COOH in *C. sativa*

To establish stable production of the pheromone precursors E/Z10-16:COOH in Camelina, we introduced a construct containing three copies of *Cpun_LATN* expression cassettes, comprising the native insect coding sequence and two codon-optimized variants driven by different promoters, along with TE16, which releases palmitic acid (16:0) from fatty acid synthesis in the chloroplasts and increases its availability to ER-bound FADs. Following seletion over four generations, we obtained *C. sativa* lines that stably accumulated E/Z10-16:COOH, without significant effects on seed weight, germination rate, or overall plant growth (Figure 5A, Figure S6).

**Figure 5.**
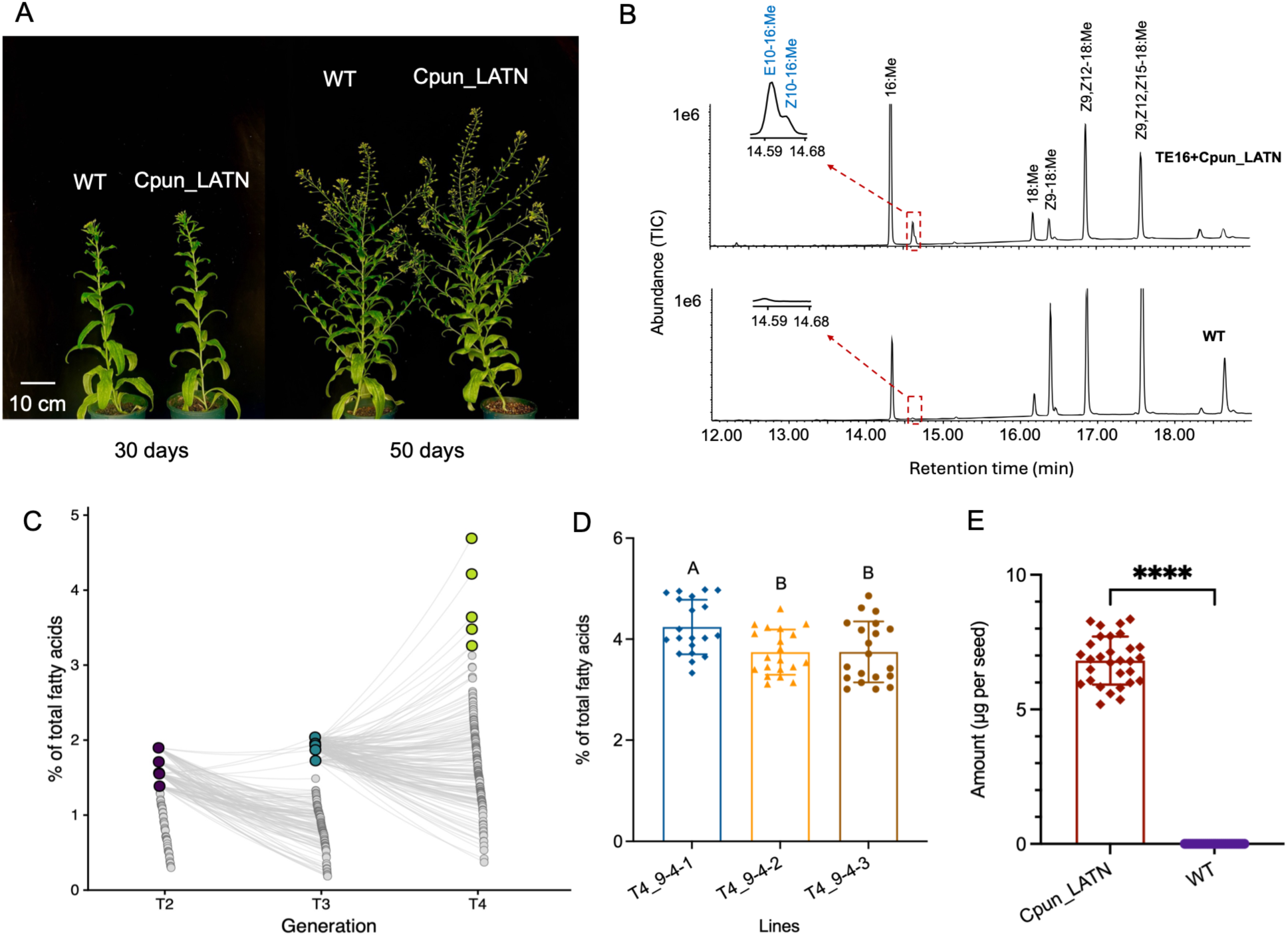
Metabolic engineering of E/Z10-16 pheromone precursor biosynthesis in *C. sativa*. (A) Phenotypes of transgenic *C. sativa* lines engineered to produce E/Z10-16 fatty acid derivatives, compared with wild-type (WT) plants. Images were taken from 30- and 50-day-old plants. Scale bar: 10 cm. (B) GC/MS analyses of fatty acids in Camelina seeds from transformant Cpun_LATN and wild type. The insets show the chromatographic peak shape details in the region where the target compounds elute. TIC: total ion current. (C) Percentage of E/Z10-16:COOH of total fatty acids in T2, T3 and T4 Camelina seeds lines. The fatty acids were analyzed in the form of corresponding methyl esters. (D) Percentage of E/Z10-16:COOH of total fatty acids in twenty individual T4 Camelina seeds from each of the top three lines with the highest target compound accumulation. Bars with the same letter are not significantly different (one-way ANOVA followed by Tukey’s HSD test). Error bars represent SEM. (E) Quantification of E/Z10-16:me in Camelina individual seeds (n=20). P value is based on t-tests:****, P < 0.0001. Error bars indicate the SEM.

GC/MS analysis of seed fatty acids showed that in wild type Camelina seeds, linolenic acid (18:3) is the most abundant fatty acid, followed by linoleic acid (18:2) and oleic acid (18:1). In constrast, the introduction of TE16 and Cpun_LATN significantly increased the accumulation of 16:0 and resulted in the production of the target E/Z10-16:COOH (Figure 5B). Progressive selection increased target fatty acid accumulation from the T2 to T4 generations, with the highest T4 lines reaching 4.7% of total fatty acids (Figure 5C). Single-seed GC/MS analysis of selected lines showed no detectable segregation of the target phenotype in the T4 generation, consistent with stable inheritance of the engineered trait (Figure 5D; Table S4). In the highest producing line, individual seeds contained an average of 6.82 μg E/Z10-16:COOH, with a maximum of 8.36 μg per seed (Figure 5E; Table S5).

## 3 Discussion

In this study, we elucidated the sex pheromone biosynthetic pathway in *C. punctiferalis*, identified a rare Δ10 desaturase and a pheromone gland fatty acyl reductase involved in this pathway, and translated this knowledge into stable bioproduction of the pheromone precursor fatty acids E10-16:COOH and Z10-16:COOH in transgenic *C. sativa* seeds. The functional reconstruction of Δ10 desaturation in an oilseed crop expands the portfolio of unusual fatty acid structures accessible through plant metabolic engineering and extends plant-based pheromone bioproduction to precursors containing an uncommon double-bond position, thereby supporting the development of biotechnological production systems for pheromone-based management of additional pests.

The incorporation of deuterium from D3-16:COOH into E/Z10-16:COOH, E/Z10-16:OH, and ultimately E/Z10-16:Ald provides direct evidence that palmitic acid is the saturated precursor of the major pheromone components. Together with the functional characterization of *Cpun_LATN* and *Cpun_3376*, these results support a biosynthetic pathway in which a Δ10 desaturase acts on palmitic acid to produce both E10- and Z10-16:COOH, which are subsequently reduced to the corresponding alcohols and finally oxidized to the aldehyde pheromone components. This pathway is consistent with the classic Type I sex pheromone biosynthetic route in moths (see e.g. Tillman et al. 1999; Jurenka 2021), but involves the comparatively unusual introduction of a double bond at the Δ10 position.

Whereas Δ9 and Δ11 desaturases have been extensively functionally characterized and are known to be key contributors of the structural diversity in moth pheromones, Δ10 desaturation remains comparatively poorly characterized (Jurenka 2021). Δ10 desaturation plays a role in extending structural diversity beyond species using Δ10-16 compounds directly, as Δ10-16 intermediates can undergo chain-shortening to generate pheromone components with double bonds at other positions. For example, in the tortricid *P. octo,* the *Δ*10 desaturase Poct_AATQ produces Z10-16:COOH which is subsequently chain-shortened to generate the Z8-14 precursor of the major pheromone component Z8-14:OAc (Hao et al 2002). This is a pathway likely followed by many other tortricid moths (Lassance et al. 2021). Interestingly, our phylogenetic analysis places Cpun_LATN within the Lepidoptera-specific so-called Δ11 desaturase clade (Lienard et al. 2008). This suggests that the Δ10 activity likely evolved within a lineage otherwise dominated by Δ11 activity. To our knowledge, the only other functionally confirmed Δ10 desaturase (Poct_AATQ; Hao et al. 2002) is from the New Zealand leafroller moth *P. octo* (Lepidoptera: Tortricidae). It should be noted that Tortricidae and Crambidae (to which *C. punctiferalis* belongs) diverged approximately 126 million years ago. Moreover, the Δ11 desaturase lineage comprises multiple paralogous copies and has undergone extensive gene duplication and loss, consistent with the birth-death evolution characteristic of insect FAD multigene families (Lassance et al. 2021; Roelofs and Rooney 2003; Helmkampf et al. 2015). Together, these observations make it unlikely to the Δ10 activities involved in pheromone biosynthesis in *P. octo* and *C. punctiferalis* represent a function inherited from a single orthologous gene. Rather, Δ10 regioselectivity may have evolved independently in different members of the lineage, representing an example of convergent evolution towards Δ10 desaturation.

The Poct_AATQ desaturase produces exclusively Z10-16:COOH from palmitic acid. In contrast, Cpun_LATN produces both the *E* and *Z* isomers, although in unequal proportions. The ability of a single desaturase to generate both geometric isomers provides the biochemical basis for production of both Δ10 components of the *C. punctiferalis* pheromone blend without requiring separate E- and Z-specific desaturase, thus reducing the enzymatic complexity required to produce the species-specific precursors. Interestingly, expression of *Cpun_LATN* in both *N. benthamiana* and *C. sativa* produced E10- and Z10-16:COOH at a ratio of approximate 3:1. This ratio is consistent with the intermediate profile observed in the insect pheromone glands (Figure 1B), indicating that Cpun_LATN exhibits similar stereoselectivity *in planta* and in its native context. In contrast, the final pheromone blend contains E10- and Z10-16:Ald at a ratio of approximately 9:1, indicating that the initial E/Z bias generated by Cpun_LATN is further modified downstream, specifically during the reduction step. Consistent with this interpretation, Cpun_3376 exhibited activity towards both E10- and Z10-16 substrates in yeast, but displayed a pronounced preference for the E isomer. Such an isomer–selective reduction step has also been documented for pheromone-biosynthetic FARs in *Ostrinia* spp. (Lepidoptera: Crambidae) (Lassance et al. 2010; 2013), supporting that FARs can play an active role in shaping pheromone blend composition rather than merely mediating conversion of fatty acid precursors to primary alcohols. Because the terminal oxidation step converting fatty alcohols to aldehydes has not yet been molecularly resolved in moths, reconstruction of the complete pheromone biosynthetic pathway in heterologous host is currently not possible using moth-derived enzymes alone. Moreover, stable accumulation of fatty acid and alcohol intermediates is likely more compatible with plant and microbial production systems than accumulation of reactive and volatile aldehydes, which could complicate extraction, storage, and downstream chemical conversion workflows. Fatty acids especially represent suitable targets for bioproduction because they can accumulate in seed lipids and be subsequently converted to the corresponding alcohol, acetate ester or aldehyde pheromones through downstream biocatalytic or chemical processes. We therefore focused the stable plant engineering on production of the E/Z10-16 fatty-acid precursors.

The successful synthesis of E/Z10-16:COOH in *N. benthamiana* confirmed the Δ10-desaturase function of *Cpun_LATN* in plants. Importantly, stable expression of *Cpun_LATN* together with the thioesterase *TE16* resulted in the accumulation of E/Z10-16:COOH in *C. sativa* seeds, demonstrating that this unusual Δ10 desaturation can be fully reconstructed in an oilseed crop. Following progressive selection over four generations, the best-performing T4 lines accumulated up to 4.7% E/Z10–16:COOH in total seed lipids, with individual seeds containing up to 8.3 μg of the target compounds. This is notable because, whereas Δ9 and Δ11 desaturases have been extensively characterized and engineered in heterologous systems (Knipple et al. 1998; Nakamura and Nara 2004; Tonon et al. 2004; Lienard et al. 2008; Ding et al. 2014; Nagao et al. 2019; Cheeld et al. 2020; Wang et al. 2022; Liu et al. 2026), desaturases introducing double bond at less common positions have often proven more difficult to functionally reconstitute in heterologous expression systems. For instance, the functional expression of the Δ8 desaturase from *Grapholita molesta* (oriental fruit moth) (Dam et al. 2025) and the Δ7 desaturase from *Lobesia botrana* (European grapevine moth) (Ding et al. 2021) has proven challenging across plant, yeast, and insect-cell expression systems. The stable production of Δ10-unsaturated fatty acids in Camelina therefore expands the range of pheromone precursor structures accessible through oilseed metabolic engineering.

Although our results demonstrate that *C. sativa* can be used as a stable platform for the production of Δ10–unsaturated pheromone precursors, the maximum accumulation of 4.7% of total seed fatty acids remains below that previously achieved for Z11–16:COOH (up to 22% in Camelina; Wang et al. 2022), while exceeding the levels reported for Z13–18:COOH (Liu et al. 2026). Still, from an applied perspective, considering that single- or two-application mating disruption against *C. punctiferalis* requires 50 or 100 g/ha, respectively (Kim et al. 2024), one hectare of our highest-yielding Cpun_LATN Camelina line could theoretically produce sufficient pheromone precursor to protect approximately 224 or 112 ha of orchard, respectively, assuming 70% recovery/conversion. Nevertheless, additional optimization will be necessary to increase the yield of E/Z10–16:COOH further.

Further improvements in E/Z10-16:COOH accumulation could involve optimization of precursor supply and its conversion by Cpun_LATN, as well as downstream incorporation of the Δ10 products into storage lipids. Palmitic acid is the immediate substrate for the production of monounsaturated C16 acids and is naturally present in Camelina seed oil. Expression of *TE16* further increased its abundance, indicating enhanced availability of this substrate. Additional increases in the 16:0-ACP pool could potentially be achieved by limiting its elongation to C18 through modulation of ß-ketoacyl-ACP synthase II (KASII) (Nguyen et al. 2015). The presence of three distinct *Cpun_LATN* expression cassettes in our construct was intended to promote high desaturase expression. However, increasing transcript abundance alone may not be sufficient to substantially improve product accumulation. Several factors could constrain Cpun_LATN activity in the heterologous plant environment. Desaturase function relies on an electron transport chain that proceeds sequentially from NADH (or NADPH) to cytochrome b5 reductase (b5R), then to cytochrome b5 (cyt b5), and finally to the desaturases (Shen et al. 2020). Suboptimal interactions between the insect desaturase and the endogenous Camelina electron-transfer partners could limit Δ10 desaturation. Co-expression of compatible electron-transfer partners therefore represents a potential strategy to improve Cpun_LATN activity. Similarly, differences between the native insect and plant endoplasmic-reticulum membrane environments could further affect desaturase activity (Bernaudat et al. 2011). Finally, accumulation of the target fatty acids may also depend on their efficient sequestration into storage lipids. Native *C. sativa* diacylglyerol acyltransferases (DGATs) may have limited activity towards lipids containing E/Z10-16:COOH, potentially restricting their incorporation into triacylglycerols (TAGs) and favoring their retention in membrane lipids or other metabolic pools (Bates et al. 2014; Reynolds et al. 2017). Engineering TAG assembly for improved utilization of Δ10-16 substrates therefore represents another potential route toward increased accumulation in seeds.

Together, our results connect the biochemical evolution of an unusual moth pheromone pathway with its reconstruction in an oilseed crop. The identification of a desaturase capable of generating both E10- and Z10-16 fatty acids precursors, combined with their stable accumulation in Camelina seeds, extends the enzymatic and metabolic toolkit available for plant-based production of structurally diverse pheromone precursors. Further optimization of precursor conversion and incorporation into seed storage lipids should provide opportunities to increase yields and advance Δ10-unsaturated fatty acids towards practical pheromone production. Yet, our study provides a foundation for developing practical, cost-effective, and environmentally sustainable pheromone production systems for the management of *C. punctiferalis*.

## 4 Methods

### 4.1 Insects and Plants

Eggs of *C. punctiferalis* were purchased from Jilin Haokang Biotechnology Company (Jiylin, China). Upon hatching, larvae were reared on fresh corn, under the conditions of 16L:8D photoperiod, 65% relative humidity and a temperature of 25 ± 1 °C. The pupae were sexed and kept separately. Adults had access to a 10% honey solution as food source.

The wild type *N. benthamiana* and *C. sativa* cv. Sunesson plants were grown in a greenhouse under 16L:8D photoperiod, 60% relative humidity and a temperature of 25 ± 1 °C, with supplemental lighting (400–540 μmol/m^2^/s) to maintain day length conditions.

### 4.2 Chemicals

[16,16,16-^2^H_3_] hexadecanoic acid (D_3_–16:COOH) used for *in vivo* labeling was purchased from Larodan AB (Malmö, Sweden) and so was (*Z*)-10-hexadecenol (Z10-16:OH). (*E*)-10-hexadecenyl acetate (E10-16:OAc) was purchased from J&K Scientific (Beijing, China) and converted to the corresponding alcohol (*E*)-10-hexadecenol (E10-16:OH) under alkaline-methanol condition. These alcohols were used to synthesize the corresponding carboxylic acids and aldehydes catalyzed by pyridinium dichromate and pyridine chlorochromate, respectively. The acids were further methylated to (*Z*)- and (*E*)-10-hexadecenoate methyl esters (Z/E10-16:Me) by acid-catalyzed methanolysis. The isomeric purities of the methyl esters were measured as 97% for Z10-16:Me (including about 3% E isomer) and >99.5% for E10-16:Me, respectively. All other chemicals (solvents: heptane, chloroform, methanol, diethyl ether, dimethylsulfoxide; dimethyl disulfide (DMDS); iodine; sodium thiosulfate) were purchased from Merck (Darmstadt, Germany) (Table S6). The amounts of E/Z10-16:Me and E/Z10-16:OH were estimated based on the dosage and peak area of the reference standards E10-16:Me and E10-16:OH, respectively (Figure S7, Table S7).

### 4.3 Pheromone Component and Precursor Analysis

For pheromone component analysis using GC/MS, pheromone glands of 2- to 3-day-old virgin females (pool of 20 glands per sample; six replicates) were dissected 1-3 h into the scotophase and extracted in 100 μL heptane. After 15 min, the extract was transferred to a new vial and concentrated to a volume of ∼30 μL under a gentle stream of N_2_.

Following volatile extraction, the tissue samples were incubated overnight in 100 μL chloroform: methanol (2:1 v/v) at room temperature for total lipid extraction and precursor analysis. Subsequently, 100 μL of double-distilled water (ddH₂O) was added, the mixture was vortexed briefly (15 seconds), and the lower chloroform layer containing the extracted lipids was transferred to a new vial. The sample was then subjected to base methanolysis for conversion of total fatty acids into their corresponding fatty acid methyl esters (FAME) following the method of Bjostad and Roelofs (1984).

Double-bond positions in monounsaturated compounds were confirmed by dimethyl disulfide (DMDS) derivatization (Buser et al. 1983). Briefly, 20-50 μL of sample was treated with 100 μL of DMDS and 20 μL 5% iodine in diethyl ether. The mixture was incubated at 40 °C overnight. Afterwards, the reaction mixture was neutralized with 50 μL of a 5% aqueous sodium thiosulfate, and the resulting DMDS-adducts were extracted with 50 μL heptane for GC/MS analysis.

### 4.4 Labeling Experiments and Sample Preparation

The deuterium-labeled precursor D_3_–16:COOH was dissolved in dimethylsulfoxide (DMSO) to a concentration of 40 μg/μL. About one hour into the scotophase, 0.2 μL of the solution was topically applied to the pheromone gland of each 2- to 3-day-old female. In parallel, an equivalent volume of DMSO was applied to a separate cohort as control group. After a 50-min incubation, pheromone glands were excised. Each sample contained a pool of 20 glands. Pheromone extracts and FAME samples were prepared as described above. Five replicates were performed for both the deuterium-labeled and control samples.

### 4.5 Gas Chromatography/Mass Spectrometry (GC/MS)

Pheromone and FAME extracts were analyzed using a 5977C mass selective detector (EI mode, 70 eV; Agilent Technologies,) coupled to an 8890 series gas chromatograph, equipped with a polar column (HP-INNOWax; 30 m × 0.25 mm; 0.25 μm film thickness) (Agilent Technologies, Santa Clara, CA, USA). The oven temperature program was as follows: 80 °C for 1 min, then increased to 230 °C at a rate of 10 °C/min, and maintained for 10 min. The injector port was kept at 250 °C and helium was used as carrier gas at a flow rate of 1 mL/min. Compounds were identified by comparison of retention times and mass spectra with corresponding authentic standards.

DMDS samples were analyzed using the same instrument equipped with a non-polar column (HP-5MS; 30 m × 0.25 mm; 0.25 μm film thickness) (Agilent Technologies, Santa Clara, CA, USA). The oven temperature was set at 80°C for 1min, raised to 140°C at a rate of 20°C/ min, then to 250 °C at a rate of 4 °C/min and held for 20 min.

Incorporation of deuterium label into the pheromone components was detected by selected ion monitoring (SIM): *m/z* 74 and 270 for 16:Me, *m/z* 74 and 273 for D_3_-16:Me; *m/z* 74, 236 and 268 for E/Z10-16:Me, *m/z* 239 and 271 for D_3_-E/Z10-16:Me; *m/z* 31, 224 and 242 for 16:OH, *m/z* 31, 227 and 245 for D_3_-16:OH; *m/z* 31, 222 and 240 for E/Z10-16:OH, *m/z* 31, 225 and 243 for D_3_-E/Z10-16:OH; *m/z* 196, 222 and 240 for 16:Ald, *m/z* 199, 225 and 243 for D_3_-16:Ald; *m/z* 194, 220 and 238 for E/Z10-16:Ald, *m/z* 197, 223 and 241 for D_3_-E/Z10-16:Ald.

### 4.6 RNA Extraction, Transcriptome Sequencing, Assembly, and Annotation

Female pheromone glands (50 glands for one sample), female antennae (20 pairs of antennae for one sample), female abdomens (3 abdomens for one sample) and male abdominal tips (50 abdominal tips for one sample) from 2- to 3-day-old virgin *C. punctiferalis* adults were collected for transcriptome sequencing. The dissection was performed under a stereomicroscope 1-3 h into the scotophase. The collected tissues were immediately frozen on dry ice and then stored at −80 °C until use. The samples were homogenized, and total RNA was extracted using Trizol reagent (Invitrogen, Carlsbad, CA, USA) following the manufacturer’s instructions. The RNA samples were sent to Metware (Wuhan, China) for Illumina sequencing after quality and quantity verification by a BioAnalyzer (Agilent). Quality of the raw read data was assessed using FastQC (http://www.bioinformatics.babraham.ac.uk/projects/fastqc/). Adaptors were trimmed, low-quality reads were filtered and contaminants were removed using Trimmomatic (v0.36). Clean reads were then *de novo* assembled using Trinity (v2.8.2) with default parameters except for -- est_method RSEM --aln method bowtie2 --trinity_mode. Initial functional annotation of unigenes was performed by blasting against the Nr (NCBI non-redundant protein sequences), KOG/COG (Clusters of Orthologous Groups of proteins) and Swiss-Prot databases with threshold E-value < 1e−5. Additional blast searches were conducted using the identified *C. punctiferalis* desaturase and reductase genes as queries and the transcriptome as a custom database to ensure that all desaturase and reductase genes were discovered. The expression levels of FAD and FAR genes were estimated using RSEM (Li and Dewey 2011) and expressed as FPKM values (fragments per kilobase of transcript per million mapped reads).

### 4.7 Phylogenetic Analysis

The phylogenetic relationships of the FADs and FARs, respectively, were built based on the predicted amino acid sequences identified from *C. punctiferalis* and functionlly characterized desaturases and reductases of other moth species retrieved from National Center for Biotechnology Information (NCBI) (https://www.ncbi.nlm.nih.gov/). The sequence alignment was performed using MAFFT (Katoh et al. 2002), and maximum-likelihood trees of two gene families were built using FastTree 2.1.11, implemented in Geneious R9 (Biomatters Ltd. Auckland, New Zealand). Sequences shorter than 200 amino acids were excluded to ensure quality of the alignment and the accuracy of the phylogenetic trees. The tree was viewed and color coded using FigTree v1.4.4.

### 4.8 Cloning of *Cpun_LATN* and *Cpun_3376* and Functional Assay in *N. benthamiana*

First-strand cDNA of pheromone glands was synthesized from 1 μg of total RNA using the SuperScript^TM^ IV First-Strand Synthesis System (Invitrogen, Carlsbad, CA, USA) following the manufacturer’s protocol. The FAD *Cpun_LATN* and the FAR *Cpun_3376* (amino acid sequences are available in the Supporting Information) were amplified using gene specific primers (Tsingke Biotechnology, Hangzhou, China). To prepare for subsequent LR reaction, additional two rounds of PCR were performed to attach attL1 and attL2 at the 5’ and 3’ ends of the candidates, respectively.

The PCR product from the previous round was used as the template for the next round. All primers used are listed in Table S8. The PCR products from the final round were analyzed on 1% TAE agarose gels and then purified using GeneJET Gel Extraction Kit (Thermo Fisher Scientific, Waltham, MA, USA). The purified gene fragments were subcloned into the plant expression vector pXZP393 using Gateway technology (Thermo Fisher). Plasmids were subjected to Sanger sequencing by the company Tsingke Biotechnology (Hangzhou, China).

The constructs containing Cpun*_LATN* and *Cpun_3376* genes were introduced into *Agrobacterium tumefaciens* GV3101 strain following a freeze-thaw transformation protocol that involves sequential incubation on ice (5 min), in liquid nitrogen (5 min), and at 37°C (5 min). P19 is a viral silencing suppressor protein that prevents the plant from degrading the viral RNA, allowing the viral replication and spread. The FatB type acyl-ACP thioesterase-1 from *Cuphea avigera* var. *pulcherrima* (*TE16*; Genbank accession number: KC675176.1) promotes palmitic acid production, providing more substrate for fatty acyl desaturases acting on C16:0. These two genes were synthesized (Tsingke Biotechnology, Hangzhou, China) and introduced into *A. tumefaciens*. Agrobacterium strains containing individual gene constructs were cultured overnight in 10 mL of LB medium with spectinomycin (75 mg/L) at 30 °C and 250 rpm. The cultures were then supplied with 100 μM acetosyringone and incubated for an additional 2-3 hours to induce the Agrobacterium virulence genes. Cells were harvested and resuspended in infiltration buffer (5 mM MgCl2, 5 mM 4-morpholineethanesulfonic acid, 100 μM acetosyringone, pH 5.7). Two distinct 20 mL Agrobacterium mixtures were prepared for the functional characterization of desaturase and reductase, respectively. The desaturase assay mixture contained *P19*-, *TE16*-, and *Cpun_LATN*-harboring Agrobacterium strains, while the reductase assay mixture contained *P19*-, *TE16*-, *Cpun_LATN*- and *Cpun_3376*-harboring Agrobacterium strains. The final concentration of each Agrobacterium culture was adjusted to an OD_600_ of 0.2.

One mL of Agrobacterium mixture was infiltrated into the abaxial side of the *N. benthamiana* leaves using a syringe. Three leaves of comparable developmental stage from five randomly selected plants were infiltrated. The infiltrated plants were then maintained in a growth chamber under well-watered conditions for 4 days, after which one healthy infiltrated leaf from each plant was collected for analysis.

For fatty acids analysis, 0.7 g of fresh leaf tissue from each sample was collected into a 4 mL vial, then 1 mL of 2 % sulfuric acid in methanol was added, and the sample was incubated for methylation at 90 °C for 1 h. Subsequently, 1 mL water and 1 mL heptane were added, and the mixture was vortexed vigorously for 15 s. Following phase separation, the upper heptane layer containing the fatty acid methyl esters (FAMEs) was transferred to a 2 mL vial for GC/MS analysis.

For fatty alcohols, 0.7 g of leaf tissue was homogenized in a glass grinder with 3.75 mL of methanol/chloroform (2:1, v/v). The crude extract was transferred to a 10 mL glass tube, and 2 mL of water was added to produce a biphasic mixture. After vigorous vortexing and centrifugation at 1000×g for 1 min, the lower chloroform layer containing the total lipids was collected into a 2 mL glass tube and then was concentrated to about 50 μL under a gentle nitrogen flow. The concentrated sample was then loaded onto a TLC plate (Silica gel 60, Merck, Germany) and separated using a mixture of hexane/diethyl ether/acetic acid (85:15:1, v/v/v). The plate was lightly sprayed with water to visualize the bands. The target bands were scraped into 4 mL vials and extracted with 1 mL of methanol/chloroform (2:1, v/v) in a sonication bath for 30s. After centrifugation at 1000×g for 1 min, the supernatant was transferred to a new tube, and 1 mL of water was added to re-extract the lipids into the chloroform. The chloroform phase containing the alcohols was collected into a new vial for GC/MS analysis.

### 4.9 Cloning of *Cpun_LATN* and *Cpun_3376* and Functional Assay in Yeast

The purified *Cpun_LATN* and *Cpun_3376* open reading frames were cloned into yeast expression vectors pYEX-CHT and pYES-DEST52 using Gateway technology (Thermo Fisher), respectively. These constructs were then transformed into the *Saccharomyces cerevisiae* yeast strains of Δ9 desaturase-deficient Δ*ole1* and INV*Sc*1, respectively, using S.c. easy yeast transformation kit (Thermo Scientific^TM^) following the manufacturer’s protocols. Positive transformants were selected at 30 °C on SC-URA plates containing 0.7% yeast nitrogen base (Sigma), 0.192% dropout medium without uracil (Formedium), 0.01% adenine (Sigma-Aldrich), 1% tergitol (Sigma-Aldrich), 2% (w/v) agar, 2% (w/v) glucose (Merck) and 0.25 mM oleic acid (Sigma-Aldrich). For functional assays, individual yeast colonies were grown in 5mL of the same selective medium at 30 °C and 300 rpm for 48h, diluted to an OD_600_ of 0.4, and then 2 mL of the diluted cultures were incubated with substrates. For Cpun_LATN, the substrates were 0.25 mM each of 12:Me, 14:Me and 16:Me, with 2 mM CuSO_4_ added for induction. For Cpun_3376, 0.25 mM E10-16:Me or Z10-16:Me was used, with 2% galactose added for induction. After 48 h, lipids from Cpun_LATN yeast cells were extracted with chloroform/methanol (2:1 v:v), subjected to acid-catalyzed methanolysis, and the resulting fatty acid methyl esters were analyzed by GC/MS. Alcohols from Cpun_3376 cultures were extracted with heptane and directly analyzed by GC/MS.

### 4.10 Construction and Screening of Transgenic *C.sativa* strain for production of E/Z10- 16:COOH

The full-length *TE16*, the native insect *Cpun_LATN* gene and two codon optimized genes of *Cpun_LATNs* (synthesized by Tsingke Biotechnology, Hangzhou, China) were assembled into the plant expression vector pBinGlyBar (Nguyen et al. 2013). Their expression was driven by four distinct seed-specific promoters. The final constructed plasmid contained four exogenous genes. *TE16* was driven by *glycinin* promotor, and the three copies of *Cpun_LATN* gene (one native copy from insects, one codon optimized from *N. benthamiana*, one codon optimized from *Arabidopsis thaliana*) were controlled by *oleosin*, *β-conglycinin* and *napin* promotors, respectively (Figure S8). The entire assembly, including all genes with their promoters and terminators, was performed using Multisite Gateway® recombination cloning technology (Invitrogen) into the pBinGlyBar vector, which contains a marker gene for Basta selection. The constructed expression clones were confirmed by Sanger sequencing (Tsingke Biotechnology, Hangzhou, China).

The construct was introduced into *A. tumefaciens* GV3101 strain as described above. A single clone from LB plate (50 mg/L Rifampicin and 50 mg/L Kanamycin) was picked and inoculated into 2mL LB medium and then scaled up to 2 L for transformation. The bacteria were harvested and resuspended in 600 mL buffer (containing 1.3 g Murishige and Skoog medium, 30 g saccharose, 300 μL Silwet77 and 100 μM acetosyringone). About 5-week-old flowering Camelina plants were transformed by floral vacuum infiltration method as previously described (Liu et al. 2012; Xia et al. 2022).

T1 seeds harvested from floral-dipped plants were sown in soil. When the plants had four leaves, herbicide was sprayed. T1 plants that survived Basta selection were considered positive transformants. T2 seeds harvested from these T1 plants were initially screened by pooling twenty seeds per line for fatty acid analysis by GC/MS. The ten T2 lines with the highest content of the target compound were advanced to single-seed fatty acid analysis, and ten individual T2 seeds per line were analyzed. The top five lines with the highest percentage of the target compound in indicidual seeds were selected, and at least twenty seeds from each line were used to produce the next generation. A similar selection procedure was conducted on the T3 and T4 seeds. Twenty individual T4 seeds from each of the top three lines with the highest content of the target compound were analysed. No segregation of the target fatty-acid phenotype was observed among T4 seeds derived from the selected T3 lines, consistent with stable inheritance of the transgene-associated phenotype and suggesting that the transgenic loci had reached homozygosity. Camelina seed samples for GC/MS were prepared by the same method as that used for *N. benthamiana*.

## Author Contributions

X.Q.H., B.J.D and C.L. conceived and designed the study. X.Q.H. and C.L. obtained funding. X.Q.H., P.H. and X.G. carried out vector construction, gene functional assays and sample analysis.

X.Q.H. and H.L.W. performed the labeling experiment. H.L.W. and Y.J. contributed to the standards and substrates preparation. B.J.D., H.L.W., and C.L. provided technical guidance and suggestions on metabolic engineering strategies. J.M.L. and X.Q.H. completed the design and standardization of figures. X.Q.H. wrote the original draft. C.L., J.M.L. and H.L.W. contributed to reviewing and editing the manuscript. All authors approved the final manuscript.

## Acknowledgements

We would like to thank Yuwan Zhao for her help with plant cultivation. This work was supported by grants from the Zhejiang Provincial Natural Science Foundation of China (ZCLZ26C1401), the Key Research and Development Program of Zhejiang Province (2024SSYS0102), and the Special Project of Xianghu Laboratory (2025X1ZN1Z010).

## Data Availability Statement

The data that support the findings of this study are available in the Supporting Information of this article. The RNA-Seq data reported in this paper have been deposited in the Genome Sequence Archive at the BIG DataCenter (http://bigd.big.ac.cn/gsa), Beijing Institute of Genomics (BIG), Chinese Academy of Sciences, under accession nos. XXXXXX.

## Supporting Information

Additional supporting information can be found online in the Supporting Information section. Figure S1: DMDS derivatization and GC-MS analysis of pheromone gland extracts. Figure S2: DMDS derivatization and GC-MS analysis of fatty acid methyl esters from pheromone gland extracts. Figure S3: Heterologous expression of desaturase candidate *Cpun_LATN* in Δ*ole1* yeast strain. Figure S4: Mass spectra of E/Z10-16:OH produced in *Nicotiana benthamiana*. Figure S5: GC/MS profile of substrates used for functional characterization of Cpun_3376 in yeast. Figure S6: Comparison of seed weight and germination rate between the Cpun_LATN T4 generation and the wild-type controls. Figure S7: Linear relationship of peak area and amount of E10-16:Me and E10-16:OH. Figure S8: Schematic representation of the expression cassettes used for pheromone precursor production in *Camelina sativa*. Table S1: List of moth species used in the FAD and FAR phylogenetic trees. Table S2: Amount of E/Z10-16:Me and E/Z10-16:OH detected from *N. benthamiana* expressing different constructs. Table S3: Amount of E/Z10-16:OH detected from yeast expressing Cpun_3376 and empty vector. Table S4: Percentage of E/Z10-16:COOH of total fatty acids in twenty individual T4 Camelina seeds from each of the top three lines with the highest target compound accumulation. Table S5: Amount (µg) of E/Z10-16:Me detected in individual seeds from the Camelina T4 line with the highest percentage content of the target compound (Cpun_LATN T4_9-4-1) and from the wild type (WT). Table S6: List of compounds used in this study, with CAS number, purity and source. Table S7: GC/MS peak area of standard E10-16:Me and E10-16:OH at 0.5 ng, 1ng, 5 ng and 10 ng. Table S8: Primers used in this study. Amino acid sequences of Cpun_LATN and Cpun_3376.

## References

Ando, T., S. Inomata, and M. Yamamoto. 2004. “Lepidopteran Sex Pheromones.” Topics in Current Chemistry 239: 51–96.

Bates, P. D., S. R. Johnson, X. Cao, J. Li, J. W. Nam, J. G. Jaworski, et al. 2014. “Fatty Acid Synthesis is Inhibited by Inefficient Utilization of Unusual Fatty Acids for Glycerolipid Assembly.” Proceedings of the National Academy of Sciences of the United States of America 111: 1204–1209.

Bernaudat, F., A. Frelet-Barrand, N. Pochon, S. Dementin, P. Hivin, S. Boutigny, et al. 2011. “Heterologous Expression of Membrane Proteins: Choosing the Appropriate Host.” PLoS One 6: e29191.

Bjostad, L. B., and W. L. Roelofs. 1983. “Sex Pheromone Biosynthesis in *Trichoplusia ni*: key Steps Involve delta-11 Desaturation and Chain-Shortening.” Science 220:1387–1389.

Bjostad, L. B., and W. L. Roelofs. 1984. “Sex Pheromone Biosynthetic Precursors in *Bombyx mori*.” Insect Biochemistry and Molecular Biology 14: 275–278.

Buser, H. R., H. Arn, P. Guerin, and S. Rauscher. 1983. “Determination of Double Bond Position in Mono-unsaturated Acetates by Mass Spectrometry of Dimethyl Disulfide Adducts.” Analytical Chemistry 55: 818–822.

Carot-Sans, G., L. Muñoz, M. D. Piulachs, A. Guerrero, and G. Rosell. 2015. “Identification and Characterization of a Fatty Acyl Reductase from a *Spodoptera littoralis* Female Gland Involved in Pheromone Biosynthesis.” Insect Molecular Biology 24: 82–92.

Chen, W. B., C. Yang, X. D. Huang, K. L. He, and Z. Y. Wang. 2022. “Screening of the Sex Pheromones Formulation and Application for Population Dynamics Monitoring of *Conogethes punctiferalis* (Guenée) in Huang-Huai-Hai Summer Corn Region in China.” Plant Protection 48: 211–219. (in Chinese)

Cheeld, H., G. Bhutada, F. Beaudoin, and P. J. Eastmond. 2020. “DES2 is a Fatty Acid Δ11 Desaturase Capable of Synthesizing Palmitvaccenic Acid in the Arbuscular Mycorrhizal Fungus *Rhizophagus Irregularis*.” FEBS Letters 594:1770–1777.

Dam, M. I., D. J. Ding, K. Brauburger, H. L. Wang, D. Powell, A. T. Groot, D. G. Heckel, and C. Löfstedt. 2025. “Sex Pheromone Biosynthesis in the Oriental Fruit Moth *Grapholita molesta* Involves Δ8 Desaturation.” Insect Biochemistry and Molecular Biology 180: 104307.

Ding, B. J., and C. Löfstedt. 2015. “Analysis of the *Agrotis segetum* Pheromone Gland Transcriptome in the Light of Sex Pheromone Biosynthesis.” BMC Genomics 16: 711.

Ding, B. J., M. A. Liénard, H. L. Wang, C. H. Zhao, and C. Löfstedt. 2011. “Terminal Fatty-acyl-CoA Desaturase Involved in Sex Pheromone Biosynthesis in the Winter Moth (*Operophtera brumata*).” Insect Biochemistry and Molecular Biology 41:715–722

Ding, B., P. Hofvander, H. Wang, T. P. Durrett, S. Stymne, and C. Löfstedt. 2014. “A Plant Factory for Moth Pheromone Production.” Nature Communications 5: 3353.

Ding, B. J., Y. H. Xia, H. L. Wang, F. Andersson, E. Hedenström, J. Gross, and C. Löfstedt. 2021. “Biosynthesis of the Sex Pheromone Component (*E*,*Z*)-7,9-Dodecadienyl Acetate in the European Grapevine Moth, *Lobesia botrana*, Involving Δ11 Desaturation and an Elusive Δ7 Desaturase.” Journal of Chemical Ecology 47: 248–264.

Dong, Z. B., X. T. Yang, and K. F. Yang. 2024. “Control Effect of Mating Disruption on Fruit Borer Pest *Conogethes punctiferalis* in Pecan.” Modern Agricultural Science and Technology. 9: 88–99. (in Chinese)

Dou, X., A. Zhang, and R. Jurenka. 2020. “Functional Identification of Fatty Acyl Reductases in Female Pheromone Gland and Tarsi of the Corn Earworm, *Helicoverpa zea*.” Insect Biochemistry and Molecular Biology 116:103260.

Foster, S. P., and W. L. Roelofs. 1996. “Sex Pheromone Biosynthesis in the Tortricid Moth, Ctenopseustis herana (Felder & Rogenhofer).” Archives of Insect Biochemistry and Physiology 33:135–147.

Hagström, Å. K. J. Albre, L. K. Tooman, A. H. Thirmawithana, J. Corcoran, C. Löfstedt, and R. D. Newcomb. 2014. “A Novel Fatty Acyl Desaturase from the Pheromone Glands of *Ctenopseustis obliquana* and *C. herana* with Specific Z5-Desaturase Activity on Myristic Acid.” Journal of Chemical Ecology 40: 63–70.

Hagström, Å. K., M. A. Liénard, A. T. Groot, E. Hedenström, and C. Löfstedt. 2012. “Semiselective Fatty Acyl Reductases from Four Heliothine Moths Influence the Specific Pheromone Composition.” PloS One 7: e37230.

Hao, G., W. Liu, M. O’Connor, and W. L. Roelofs. 2002. “Acyl-CoA Z9-and Z10-Desaturase Genes from a New Zealand Leafroller Moth Species, *Planotortrix octo*.” Insect Biochemistry and Molecular Biology 32:961–966.

Helmkampf, M., E. Cash, and J. Gadau. 2015. “Evolution of the Insect Desaturase Gene Family with an Emphasis on Social Hymenoptera.” Molecular Biology and Evolution, 32: 456–471.

Jurenka, R. A. 2021. “Lepidoptera: Female Sex Pheromone Biosynthesis and Its Hormonal Regulation.” In Insect Pheromone Biochemistry and Molecular Biology, edited by G. J. Blomquist and R. G. Vogt, 13–88. Academic Press.

Katoh, K., K. Misawa, K. Kuma, and T. Miyata. 2002. “MAFFT: a Novel Method for Rapid Multiple Sequence Alignment Based on Fast Fourier Transform.” Nucleic Acids Research 30:3059–3066.

Kim, J., S. Jung, and Y. U. Kim. 2024. “Pheromone-Based Mating Disruption of *Conogethes punctiferalis* (Lepidoptera: Crambidae) in Chestnut Orchards.” Insects 15: 445.

Knipple, D. C., C. L. Rosenfield, R. Nielsen, K. M. You, and S. E. Jeong. 2002. “Evolution of the Integral Membrane Desaturase Gene Family in Moths and Flies.” Genetics 162: 1737–1752.

Knipple, D. C., C. L. Rosenfield, S. J. Miller, W. Liu, J. Tang, P. W. Ma, and W. L. Roelofs. 1998. “Cloning and Functional Expression of a cDNA Encoding a Pheromone Gland-specific Acyl-CoA Δ11-Desaturase of the Cabbage Looper Moth, *Trichoplusia ni*.” Proceedings of the National Academy of Sciences of the United States of America 95:15287–15292.

Konno, Y., K. Arai, K. Sekiguchi, and Y. Matsumoto. 1982. “(*E*)-10-Hexadecenal, a Sex Pheromone Component of the Yellow Peach Moth, *Dichocrocis punctiferalis* Guenée (Lepidoptera: Pyralidae).” Applied Entomology and Zoology 17: 207–217.

Lassance, J., B. Ding, and C. Löfstedt. 2021. “Evolution of the Codling Moth Pheromone via an Ancient Gene Duplication.” BMC Biology 19: 83.

Lassance, J. M., A. T. Groot, M. A. Liénard, B. Antony, C. Borgwardt, F. Andersson, E. Hedenström, D. G. Heckel, and C. Löfstedt. 2010. “Allelic Variation in a Fatty-acyl Reductase Gene Causes Divergence in Moth Sex Pheromones.” Nature 466: 486–489.

Lassance, J. M., M. A. Liénard, B. Antonya, S. Qian, T. Fujii, J. Tabata, et al. 2013. “Functional Consequences of Sequence Variation in the Pheromone Biosynthetic Gene pgFAR for Ostrinia Moths.” Proceedings of the National Academy of Sciences of the United States of America 110: 39673972.

Li, B., and C. N. Dewey. 2011. “RSEM: Accurate Transcript Quantification from RNA-Seq Data with or without a Reference Genome.” BMC Bioinformatics 12: 323.

Liu, M., H. Li, H. Jin, Y. Xiao, P. He, C. Tan, H. L. Wang, J. M. Lassance, C. Löfstedt and B. J. Ding. 2026. “Paving the Way for Plant-Based Bioproduction of (*Z*)-13-Octadecenoic Moth Pheromone Compounds.” Plant Biotechnology Journal 0:1–16.

Liu, M., Y. Tian, and Y. Li. 1994. “Identification of Minor Components of the Sex Pheromone of Yellow Peach Moth *Dichorocis punctifeealis* Guenée and Field Trials.” Insect Science 1: 150–155.

Liu, W., H. Jiao, N. C. Murray, M. O’Connor, and W. L. Roelofs. 2002. “Gene Characterized for Membrane Desaturase that Produces (*E*)-11 Isomers of Mono-and Diunsaturated Fatty Acids.” Proceedings of the National Academy of Sciences of the United States of America 99: 620–624.

Liu, W., A. P. Rooney, B. Xue, W. L. Roelofs. 2004. “Desaturases from the Spotted Fireworm Moth (*Choristoneura parallela*) Shed Light on the Evolutionary Origins of Novel Moth Sex Pheromone Desaturases.” Gene 342: 303– 311.

Liu, X., J. Brost, C. Hutcheon, R. Guilfoil, A. K. Wilson, S. Leung, and J. De Rocher. 2012. “Transformation of the Oilseed Crop *Camelina sativa* by *Agrobacterium*-Mediated Floral Dip and Simple Large-Scale Screening of Transformants.” In Vitro Cellular & Developmental Biology-Plant 48: 462–468.

Liénard, M. A., J. M. Lassance, H. L. Wang, C. H. Zhao, J. Piskur, T. Johansson, and C. Löfstedt. 2010. “Elucidation of the Sex-pheromone Biosynthesis Producing 5,7-Dodecadienes in *Dendrolimus punctatus* (Lepidoptera: Lasiocampidae) Reveals Δ11-and Δ9-Desaturases with Unusual Catalytic Properties.” Insect Biochemistry and Molecular Biology 40: 440–452.

Liénard, M. A., M. Strandh, E. Hedenström, T. Johansson, and C. Löfstedt. 2008. “Key Biosynthetic Gene Subfamily Recruited for Pheromone Production Prior to the Extensive Radiation of Lepidoptera.” BMC Evolutionary Biology 8: 270.

Löfstedt, C., N. Wahlberg, and J. G. Millar. 2016. “Evolutionary Patterns of Pheromone Diversity in Lepidoptera.” In Pheromone Communication in Moths: Evolution, Behavior and Application, Edited by J. D. Allison and R. T. Cardé, 43–78. University of California Press.

Löfstedt, C., and Y. Xia. 2021. “Biological Production of Insect Pheromones in Cell and Plant Factories.” In Insect Pheromone Biochemistry and Molecular Biology, edited by G. J. Blomquist and R. G. Vogt, 89–121. Academic Press.

Ma, P. W. K., and S. B. Ramaswamy. 2003. “Biology and Ultrastructure of Sex Pheromone Producing Tissue.” In Insect Pheromone Biochemistry and Molecular Biology, Edited by G. J. Blomquist and R. G. Vogt, 19–51. Academic Press.

Nagao, K., A. Murakami, and M. Umeda. 2019. “Structure and Function of Δ9-fatty Acid Desaturase.” Chemical and Pharmaceutical Bulletin 67: 327–332.

Nakamura, M. T., and T. Y. Nara. 2004. “Structure, Function, and Dietary Regulation of Delta6, Delta5, and Delta9 Desaturases.” Annual Review of Nutrition 24: 345–376.

Nam, I., J. H. Park, J. W. Lee, D. H. Lee, Jung, W. Lee. 2022. “Field confirmation of (*Z*)-9-heptacosene and (3*Z*,6*Z*,9*Z*)-tricosatriene as key sex pheromone components of Korean *Conogethes punctiferalis* Guene (Lepidoptera: Crambidae).” Journal of Economic Entomology 115: 1976–1986.

Nguyen, H. T., H. Park, K. L. Koster, R. E. Cahoon, H. T. M. Nguyen, J. Shanklin, et al. 2015. “Redirection of Metabolic Flux for High Levels of Omega-7 Monounsaturated Fatty Acid Accumulation in Camelina Seeds.” Plant Biotechnology Journal 13: 38–50.

Nguyen, H. T., J. E. Silva, R. Podicheti, J. Macrander, W. Yang, T. J. Nazarenus, et al. 2013. “Camelina Seed Transcriptome: a Tool for Meal and Oil Improvement and Translational Research.” Plant Biotechnology Journal 11: 759–769.

Reynolds, K. B., M. C. Taylor, D. P. Cullerne, C. L. Blanchard, C. C. Wood, S. P. Singh, et al. 2017. “A Reconfigured Kennedy Pathway which Promotes Efficient Accumulation of Medium-Chain Fatty Acids in Leaf Oils.” Plant Biotechnology Journal 15: 1397–1408.

Roelofs, W. L., W. Liu, G. Hao, H. Jiao, A. P. Rooney, and C. E. Linn. 2002. “Evolution of Moth Sex Pheromones via Ancestral Genes.” Proceedings of the National Academy of Sciences of the United States of America 99:13621– 13626.

Roelofs, W. L., and A. P. Rooney. 2003. “Molecular Genetics and Evolution of Pheromone Biosynthesis in Lepidoptera.” Proceedings of the National Academy of Sciences of the United States of America 100: 9179–9184.

Shen, J., G. Wu, A. L. Tsai, and M. Zhou. 2020. “Structure and Mechanism of a Unique Diiron Center in Mammalian Stearoyl-CoA Desaturase.” Journal of Molecular Biology 432: 5152–5161.

Thyagaraj, N. E., P. K. Singh, and A. K. Singh. 2003. “Bioecology of Cardamom Shoot and Fruit Borer, *Conogethes punctiferalis* Guenée.” Currents in Research 32: 3–4.

Tillman, J. A., S. J. Seybold, R. A. Jurenka, and G. J. Blomquist. 1999. “Insect Pheromones—an Overview of Biosynthesis and Endocrine Regulation.” Insect Biochemistry and Molecular Biology 29: 481–514.

Tonon, T., D. Harvey, R. Qing, Y. Li, T. R. Larson, and I. A. Graham. 2004. “Identification of a Fatty Acid Δ11-Desaturase from the Microalga *Thalassiosira pseudonana*.” FEBS Letters 563: 28–34.

Wang, H. L., B. J. Ding, J. Q. Dai, T. J. Nazarenus, R. Borges, et al. 2022. “Insect Pest Management with Sex Pheromone Precursors from Engineered Oilseed Plants.” Nature Sustainability 5: 981–990.

Wang, H. L., M. A. Liénard, C. H. Zhao, C. Z. Wang, and C. Löfstedt. 2010. “Neofunctionalization in an Ancestral Insect Desaturase Lineage Led to Rare Δ6 Pheromone Signals in the Chinese Tussah Silkworm.” Insect Biochemistry and Molecular Biology 40: 742–751.

Xia, Y. H. 2020. “Plants as Factories for Insect Biosynthetic Pathway of Female Moths.” Media-Tryck, Lund. PhD Thesis.

Xia, Y., H. Wang, B. Ding, et al. 2021. “Green Chemistry Production of Codlemone, the Sex Pheromone of the Codling Moth (*Cydia pomonella*), by Metabolic Engineering of the Oilseed Crop Camelina (*Camelina sativa*).” Journal of Chemical Ecology 47: 950–967.

Xiao, W., S. Matsuyama, T. Ando, J. G. Millar, and H. Honda. 2012. “Unsaturated Cuticular Hydrocarbons Synergize Responses to Sex Attractant Pheromone in the Yellow Peach Moth, *Conogethes punctiferalis*.” Journal of Chemical Ecology 38: 1143–1150.

