## supplemental Figures and tables for "Reconstruction of a rare Δ10 desaturation pathway enables production of yellow peach moth pheromone precursors in *Camelina sativa*"

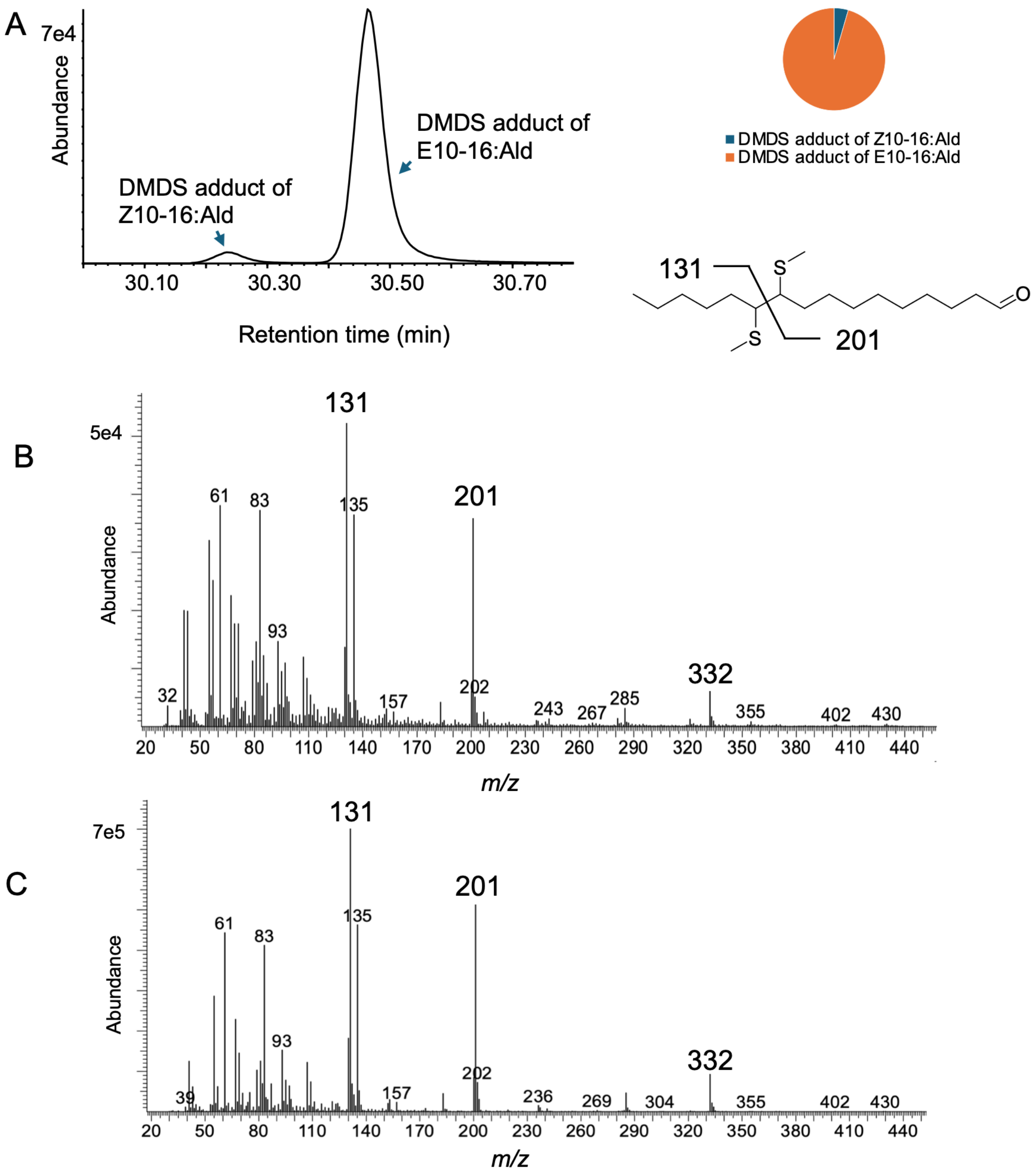


**Figure S1.** DMDS derivatization and GC-MS analysis of pheromone gland extracts. (A) Representative extracted ion chromatogram (EIC) showing the DMDS adducts of Z10-16:Ald and E10-16:Ald (*m/z*=201; left), the pie chart representing the relative chromatographic peak areas of the two DMDS adducts (upper right*)* schematic illustrating the fragmentation pattern and the origin of diagnostic ions of the DMDS adducts (lower right). (B, C) Mass spectrua of DMDS adduct of Z10-16:Ald (B) and E10-16:Ald (C). Diagnostic fragment ions (*m/z* 131 and 201) and the molecular ion (*m/z* 332) are highlighted in larger font .

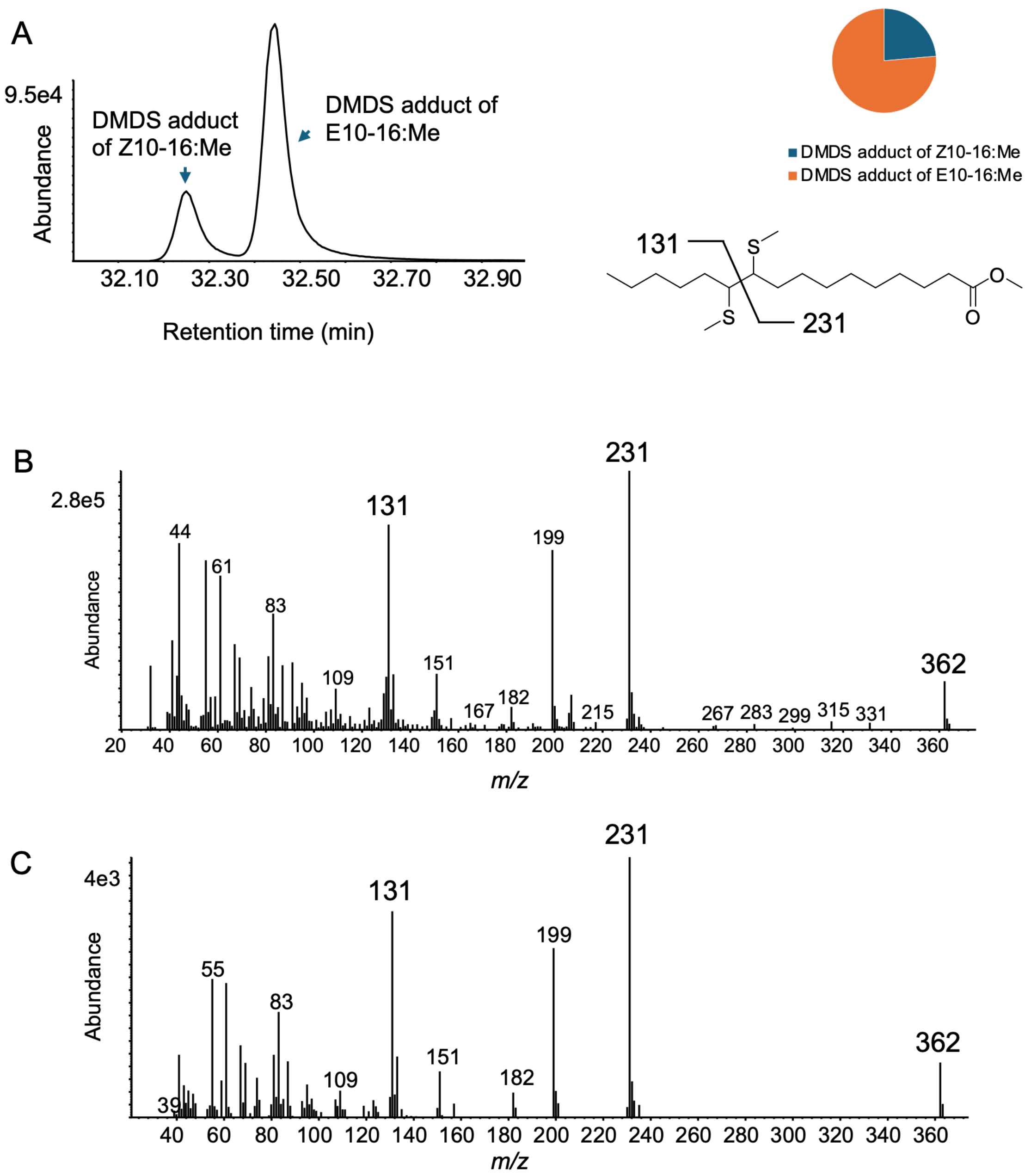


**Figure S2.** DMDS derivatization and GC-MS analysis of fatty acid methyl esters from pheromone gland extracts. (A) Representative extracted ion chromatogram (EIC) showing the abundance of DMDS adducts of Z10-16:Me and E10-16:Me (*m/z*=231; left), their relative chromatographic peak areas (upper-right), and a schematic illustrating the origin of diagnostic ions (lower right). (B, C) Mass spectra of the DMDS adduct of Z10-16:Me (B) and E10-16:Me (C). Diagnostic fragment ions (*m/z* 131 and 231) and the molecular ion (*m/z* 362) are highlighted in larger font.

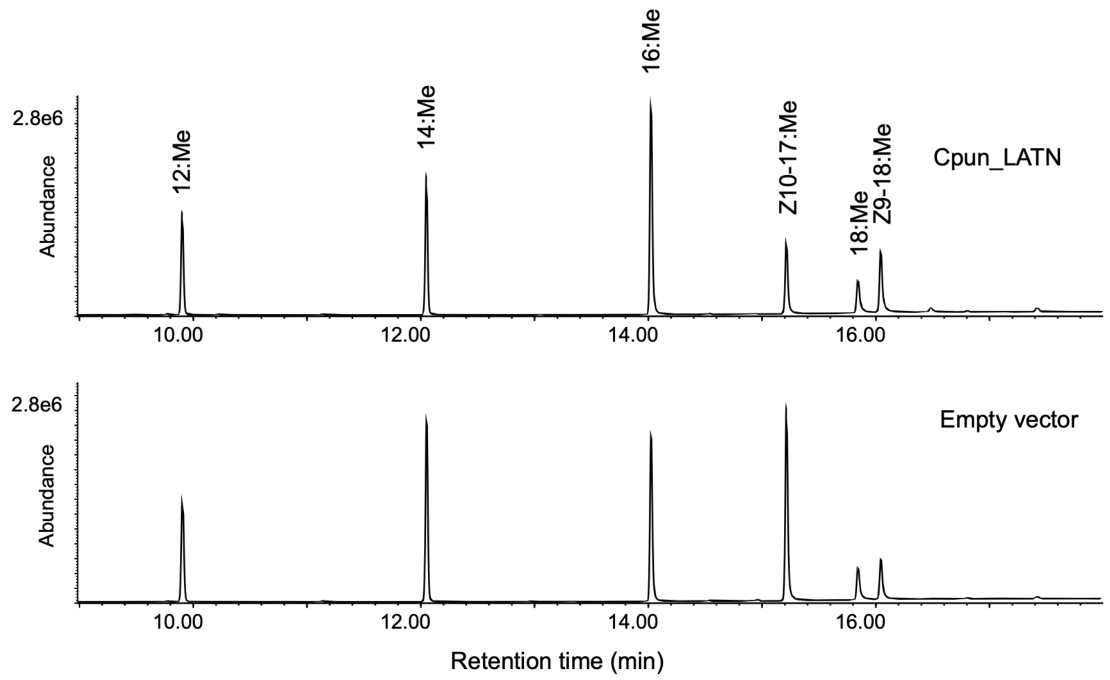


**Figure S3.** Heterologous expression of desaturase candidate Cpun_LATN in *Δole1* yeast strain. GC/MS analysis of fatty acids from yeast expressing Cpun_LATN with 12:Me, 14:Me and 16:Me added as substrates, compared with the empty vector control. Displayed chromatograms are representative examples of at least six replicates.

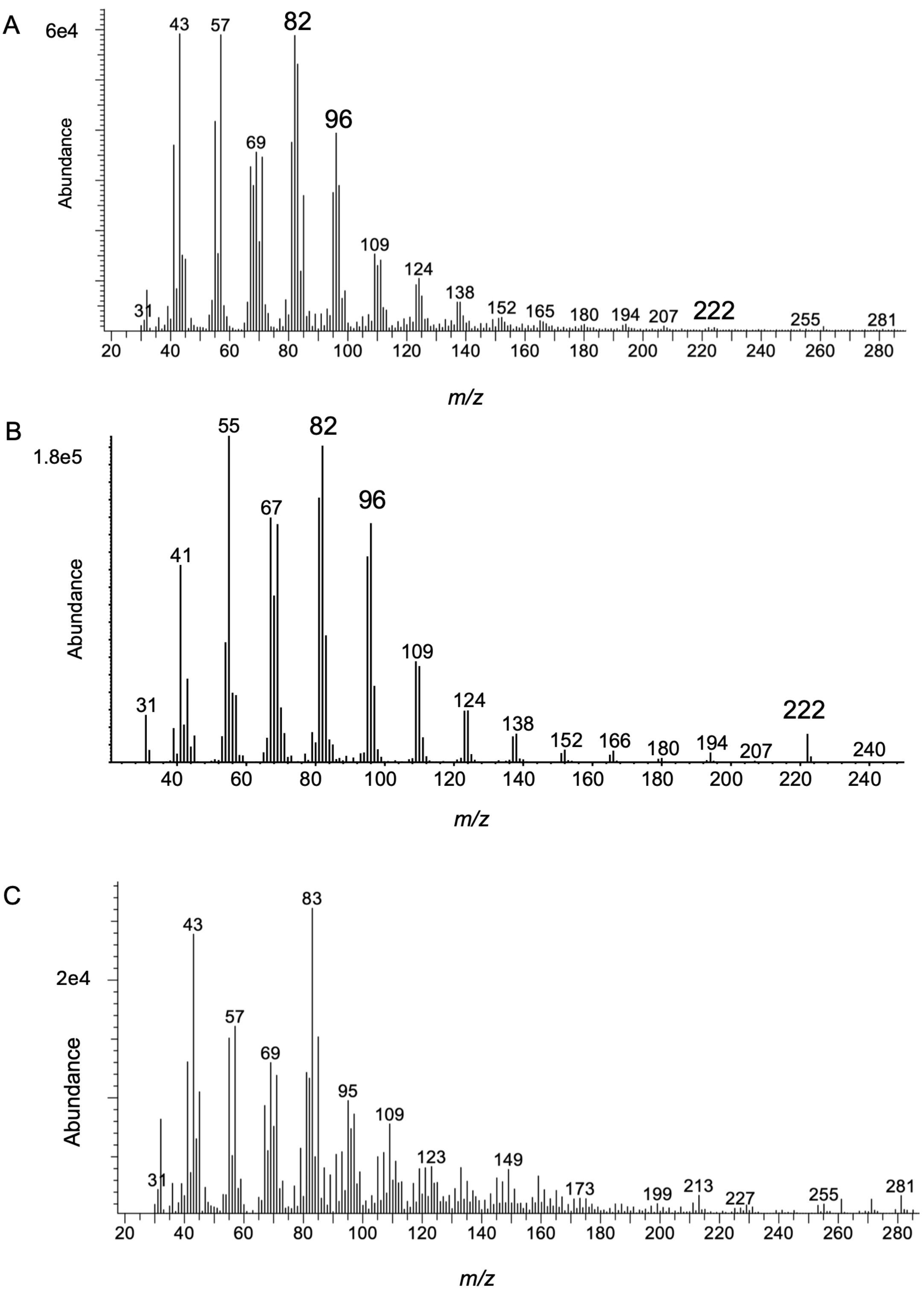


**Figure S4.** Mass spectra of E/Z10-16:OH produced in *Nicotiana benthamiana.* (A) Mass spectrum of E/Z10-16:OH detected in *N. benthamiana* co-expressing Cpun_3376, Cpun_LATN and TE16, showing a fragmentation pattern consistent with standard E10-16:OH (B). (C) Mass spectrum of the minor peak detected at the corresponding retention time in the control, showing a fragmentation pattern inconsistent with standard E10-16:OH (B).

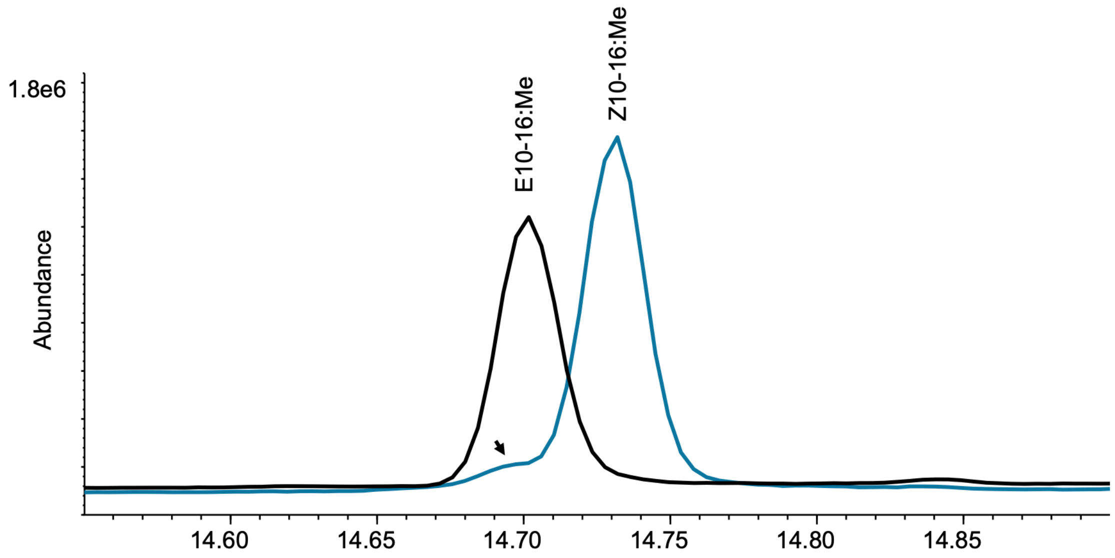


**Figure S5.** GC/MS profile of substrates used for functional characterization of Cpun_3376 in yeast. The *Z* isomer substrate contains a small amount of the corresponding *E* isomer (indicated by a black arrow).

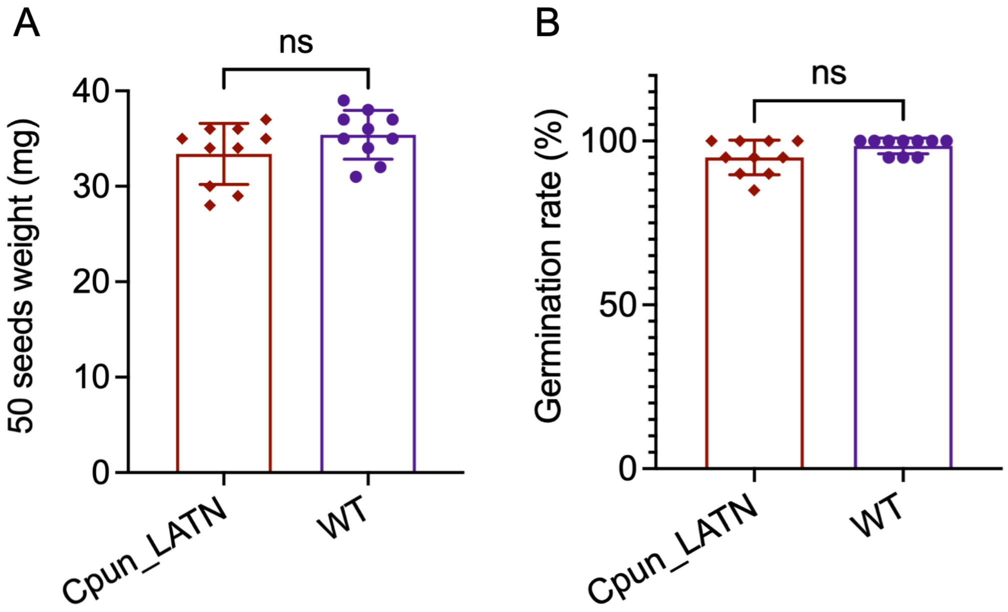


**Figure S6.** Comparison of seed weight and germination rate between the Cpun_LATN T4 generation and the wild-type controls. (A) Weight of 50 seed (n=10 replicates). (B) Seed germination rate. Bars represent means ± SEM, with individual points shown. Statistical significance was assessed using Student's t-tests (ns, not significant; $P > 0.05$).

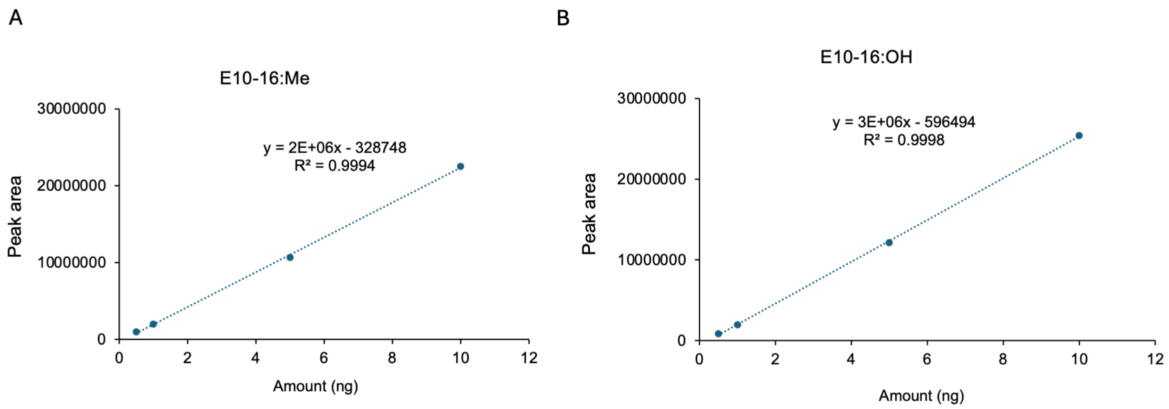


**Figure S7.** Linear relationship of peak area and amount of (A) E10-16:Me and (B) E10-16:OH.

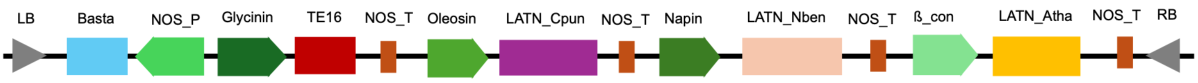


**Figure S8.** Schematic representation of the expression cassettes used for pheromone precursor production in *C. sativa.* The construct contains three versions of Cpun_LATN (the native insect coding sequence plus two versions codon-optimized for *Nicotiana benthamiana* and *Arabidopsis thaliana*) together with TE16, a gene encoding a FatB-type acyl-ACP thioesterase-1 from *Cuphea avigera* var. *pulcherrima*. LB and RB, left and right T-DNA borders, respectively; Basta, herbicide resistance selection marker; NOS_P and NOS_T, *Agrobacterium tumefaciens* nopaline synthase promoter and terminator, respectively; Glycinin, *Glycine max* glycinin promoter; Oleosin, *Brassica napus* oleosin promoter; Napin, *Brassica napus* napin promoter; ß_con, *Glycine max* ß_conglycinin promoter, Cpun_LATN_Nben and Cpun_LATN_Ath, versions of Cpun_LATN codon-optimized for *N. benthamiana* and *A. thaliana*, respectively.

**Table S1**. List of moth species used in the FAD and FAR phylogenetic trees.

| Abbreviations | Species |
|---|---|
| Aseg | *Agrotis segetum* |
| Atra | *Amyelois transitella* |
| Bmor | *Bombyx mori* |
| Cpom | *Cydia pomonella* |
| Cpun | *Conogethes punctiferalis* |
| Ecau | *Ephestia cautella* |
| Harm | *Helicoverpa armigera* |
| Hass | *Helicoverpa assulta* |
| Hsal | *Harpegnathos saltator* |
| Hvir | *Heliothis virescens* |

| | |
|---|---|
| Lbot | *Lobesia botrana* |
| Onub_E | *Ostrinia nubilalis* E strain |
| Onub_Z | *Ostrinia nubilalis* Z strain |
| Poct | *Planotortrix octo* |
| Pxyl | *Plutella xylostella* |
| Sexi | *Spodoptera exigua* |
| Sins | *Streltzoviella insularis* |
| Ycag | *Yponomeuta cagnagella* |
| Yevo | *Yponomeuta evonymella* |
| Ypad | *Yponomeuta padella* |
| Yror | *Yponomeuta rorrella* |

**Table S2**. Amount of E/Z10-16:Me and E/Z10-16:OH detected from *N. benthamiana* expressing different constructs.

| | Amount (μg) | | | |
|---|---|---|---|---|
| | E/Z10-16:Me | | E/Z10-16:OH | |
| | TE16/Cpun_LATN | TE16 | TE16/Cpun_LATN/Cpun_3376 | TE16/Cpun_LATN |
| Sample 1 | 10.11 | 0.38 | 0.47 | 0.24 |
| Sample 2 | 9.24 | 0.41 | 0.41 | 0.23 |
| Sample 3 | 6.94 | 0.49 | 0.38 | 0.24 |
| Sample 4 | 8.38 | 0.43 | 0.52 | 0.22 |
| Sample 5 | 7.05 | 0.47 | 0.38 | 0.22 |

**Table S3**. Amount of E/Z10-16:OH detected from yeast expressing Cpun_3376 and empty vector.

| | | Amount (μg) | | | |
|---|---|---|---|---|---|
| | | E10-16:OH | | Z10-16:OH | |
| Substrate added | Smples | Cpun_3376 | Empty vector | Cpun_3376 | Empty vector |
| E10-16:Me | Sample 1 | 4.59 | 0.24 | 0 | 0 |
| | Sample 2 | 3.87 | 0.26 | 0 | 0 |
| | Sample 3 | 3.04 | 0.26 | 0 | 0 |
| | Sample 4 | 2.81 | 0.25 | 0 | 0 |
| | Sample 5 | 3.56 | 0.26 | 0 | 0 |
| | Sample 6 | 2.45 | 0.26 | 0 | 0 |
| Z10-16Me | Sample 1 | 0.42 | 0.25 | 0.49 | 0 |
| | Sample 2 | 0.44 | 0.22 | 0.53 | 0 |
| | Sample 3 | 0.40 | 0.23 | 0.46 | 0 |
| | Sample 4 | 0.41 | 0.22 | 0.45 | 0 |
| | Sample 5 | 0.42 | 0.22 | 0.47 | 0 |
| | Sample 6 | 0.41 | 0.21 | 0.51 | 0 |
| No substrate | Sample 1 | 0.24 | 0.22 | 0 | 0 |
| | Sample 2 | 0.25 | 0.23 | 0 | 0 |
| | Sample 3 | 0.25 | 0.21 | 0 | 0 |
| | Sample 4 | 0.25 | 0.21 | 0 | 0 |
| | Sample 5 | 0.24 | 0.21 | 0 | 0 |
| | Sample 6 | 0.25 | 0.21 | 0 | 0 |

**Table S4**. Percentage of E/Z10-16:COOH of total fatty acids in twenty individual T4 Camelina seeds from each of the top three lines with the highest target compound accumulation.

| Samples | Cpun_LATN T4 lines | | |
|---|---|---|---|
| | T4_9-4-1 | T4_9-4-2 | T4_9-4-3 |
| 1 | 4.07 | 4.09 | 4.2 |
| 2 | 4.64 | 3.59 | 4.55 |
| 3 | 4.02 | 4.11 | 3.01 |
| 4 | 4.92 | 4.6 | 4.86 |
| 5 | 3.99 | 4.23 | 3.45 |
| 6 | 4.57 | 3.4 | 3.23 |
| 7 | 4.97 | 4.21 | 3.01 |
| 8 | 3.88 | 3.81 | 3.32 |
| 9 | 4.85 | 3.64 | 3.25 |
| 10 | 3.65 | 4.29 | 4.32 |
| 11 | 4.2 | 3.11 | 4.18 |
| 12 | 3.72 | 3.26 | 4.31 |
| 13 | 4.02 | 3.25 | 3.42 |
| 14 | 4.98 | 3.45 | 3.71 |
| 15 | 3.71 | 3.14 | 3.14 |
| 16 | 4.02 | 3.54 | 3.04 |
| 17 | 3.55 | 3.45 | 3.27 |
| 18 | 3.33 | 4.3 | 4.62 |
| 19 | 4.95 | 3.4 | 3.92 |
| 20 | 4.79 | 3.96 | 4.11 |

**Table S5**. Amount (µg) of E/Z10-16:Me detected in individual seeds from the Camelina T4 line with the highest percentage content of the target compound (Cpun_LATN T4_9-4-1) and from the wild type (WT).

| Samples | Cpun_LATN | WT |
|---|---|---|
| 1 | 6.79 | 0 |
| 2 | 6.06 | 0 |
| 3 | 7.04 | 0 |
| 4 | 5.59 | 0 |
| 5 | 6.8 | 0 |
| 6 | 7.71 | 0 |
| 7 | 7.42 | 0 |
| 8 | 7.13 | 0 |
| 9 | 8.27 | 0 |
| 10 | 6.49 | 0 |
| 11 | 6.35 | 0 |
| 12 | 6.74 | 0 |
| 13 | 7.72 | 0 |
| 14 | 7.32 | 0 |
| 15 | 6.4 | 0 |
| 16 | 6.96 | 0 |
| 17 | 5.84 | 0 |
| 18 | 7.26 | 0 |
| 19 | 6.86 | 0 |
| 20 | 5.37 | 0 |
| 21 | 5.94 | 0 |

| | | |
|---|---|---|
| 22 | 6.08 | 0 |
| 23 | 5.99 | 0 |
| 24 | 7.81 | 0 |
| 25 | 8.36 | 0 |
| 27 | 5.18 | 0 |
| 28 | 5.81 | 0 |
| 29 | 8.12 | 0 |
| 30 | 6.93 | 0 |

**Table S6.** List of compounds used in this study, with CAS number, purity and source.

| Compound | CAS number | Purity (%) | Source* |
|---|---|---|---|
| Heptane | 142-82-5 | ≥ 99 | Sigma |
| Chloroform | 67-66-3 | ≥ 99.8 | Sigma-Aldrich |
| Methanol | 67-56-1 | ≥ 99.9 | Macklin |
| Diethyl ether | 60-29-7 | ≥ 99.7 | Sinopharm Chemical Reagent Co., Ltd |
| Dimethyl sulfoxide | 67-68-5 | ≥ 99.5 | Solarbio |
| Dimethyl disulfide | 624-92-0 | ≥ 99 | J&K Scientific |
| Iodine | 7553-56-2 | ≥ 99 | Sigma |
| Sodium thiosulfate | 7772-98-7 | ≥ 99.5 | Sigma-Aldrich |

**Table S7**. GC/MS peak area of standard E10-16:Me and E10-16:OH at 0.5 ng, 1ng, 5 ng and 10 ng.

| | | Peak area | |
|---|---|---|---|
| | | E10-16:Me | E10-16:OH |
| 0.5ng | Sample 1 | 986922 | 811016 |
| | Sample 2 | 875165 | 782981 |
| | Sample 3 | 992656 | 887640 |
| | Sample 4 | 1061108 | 916745 |
| 1 ng | Sample 1 | 2005878 | 1559749 |
| | Sample 2 | 1939826 | 1801713 |
| | Sample 3 | 1928998 | 2120507 |
| | Sample 4 | 1949196 | 2356435 |
| 5 ng | Sample 1 | 10520200 | 11897299 |
| | Sample 2 | 10552187 | 12230192 |
| | Sample 3 | 10686063 | 11829574 |
| | Sample 4 | 10835463 | 12544957 |
| 10 ng | Sample 1 | 22006876 | 24324087 |
| | Sample 2 | 22447566 | 24175838 |
| | Sample 3 | 22533751 | 25374376 |
| | Sample 4 | 23083845 | 27766294 |

**Table S8**. Primers used in this study.

| Genes | Primer sequence (5'-3') |
|---|---|

| | |
|---|---|
| Cpun_LATN_F | ATGCCCCCTAACGCTGGT GACATTCAGAAG |
| Cpun_LATN_R | TTATAGTGTTTTCGGAGCTTGATCTCGACCCCACAAAT |
| Cpun_LATN_attL1 | AAATTGATGAGCAATGCTTTTTTATAATGCCAACTTTGTACAAAAAAGCAGATGCCCCCTAAC |
| Cpun_LATN_attL2 | AAATTGATAAGCAATGCTTTCTTATAATGCCAACTTTGTACAAGAAAGCTGTTATAGTGTTTTCGG |
| Cpun_3376_F | TACACATAAGAATATCGTACTAAATTGAAATAAATT |
| Cpun_3376_R | TACTTAGAATAATACTTAGTTTGATAACATTCATGT |
| Cpun_3376_attL1 | ACAAATTGATGAGCAATGCTTTTTTATAATGCCAACTTTGTACAAAAAAGCAGTACACATAAGAATAT CGTACTAAATT |
| Cpun_3376_attL2 | TGCAACAAATTGATAAGCAATGCTTTCTTATAATGCCAACTTTGTACAAGAAAGCTGCATACTTAGAA TAATACTTAGT |
| attL1 | CAAATAATGATTTTATTTTGACTGATAGTGACCTGTTCGTTGCAACAAATTGATGAGCAATG |
| attL2 | CAAATAATGATTTTATTTTGACTGATAGTGACCTGTTCGTTGCAACAAATTGATAAGCAATGCT |
| 393a_F | ACCCTTCCTCTATATAAGGAAGTTC |
| 393a_R | GCACAACAGAATTGAAAGCAAATATC |
| pBinGlyBar_F | ATTCACCTTCCTCTCTTCCCTATAAATA |
| pBinGlyBar_R | TCGTGACTCATTATAGAGAGTACTTAC |

### Predicted amino acid sequences of Cpun_LATN and Cpun_3376

>Cpun_LATN
MPPNAGDIQKALLASPLIDGHKPSRAAPRAHALSYLTIAAFTVYHLTGLYGLYLCFTSAKAAT
LVFAFFTFVAAQFGVTAGAHRLWSHNSFKAKAPLQILLMVLQTLAFQTSVAHWVRDHRLHH
QYADSDGDPHNASRGFFYSHIGWLLTAKTKDLLERGKKIDMSDIHSNPVLCFQEKYYMVLV
AGICFALPTAAPVLLWGEAPAVAWCVNMLRHVVNLQTTFMANSICHIFGYRPYDKNA**LATN**
NLYINLLILGDGFHNFHHAFPWDYQSSEWGNRNFNFTTIFIDFFAKIGWAYDRKTVSDDIVKH
RKIRAGDGTDLWGRDQAPKTL

>Cpun_3376
MDSTANTYEEIIEPSFSSSQIVQFYEGRSIFITGATGFLGTVLIEKLLFTCQNIKNIFVLMKQKRG
KTIERRLADMLESQAFHRIKRQRPDSIIKIIPIGGDLIKHNLGMTKSGIDLIQNEVSIVFHVAANI
SFNKDLHDTININTKATEKLINICKNMPHLAAFVYVSSGFSQSQRLIVDETLYDNTVPVETVYE
LLEMPKSQKLAVNLLEGRPNVYAYSKALAEEIINKYADQIPAAIVRPTLIISAMQEPLPGWITS
WAGPTAIVEAAGRGLLMCWNSDAANVIDLVPVDYVANLIIATAWDTHNRRVKNDMQGDK
RIKVFNYSSCTQNPIESGKFLSTCLELNKKTCGKDLFYSNQLFIFKNHYVGYWIVSTLFNILLT
GISIVIDMVLWFLGQNIKMFKIQRNLAKLTLAVVPFTQRQILFLNGNVRQLYKTLNSADQKLF
DFDVSHIQWENYLRDYVVGMKKYNEPFSKKF
